# Degron-controlled protein degradation in *Escherichia coli*: New Approaches and Parameters

**DOI:** 10.1101/2023.11.08.566101

**Authors:** Glen E. Cronan, Andrei Kuzminov

**Author notes:** For correspondence: Glen Cronan, B103 C&LSL, 601 South Goodwin Ave., Urbana IL 61801-3709, USA.

## Abstract

Protein degron tags have proven uniquely useful for characterization of gene function. Degrons mediate quick depletion, usually within minutes, of a protein of interest – allowing researchers to characterize cellular responses to the loss of function. To develop a general purpose degron tool in *E. coli,* we sought to build upon a previously characterized system of SspB-dependent inducible protein degradation. For this, we created a family of expression vectors containing a destabilized allele of SspB, capable of a rapid and nearly perfect “off-to-on” induction response. Using this system, we demonstrated control over several enzymes of DNA metabolism, but also found with other substates apparent limitations of a SspB-dependent system. Several degron target proteins were degraded too slowly to affect their complete depletion during active growth, whereas others appeared completely refractory to degron-promoted degradation. We demonstrated that a model substrate, beta-galactosidase, was positively recognized as a degron substrate, but failed to be degraded by the ClpXP protease — demonstrating an apparently unknown mechanism of protease resistance. Thus, only a minority of our, admittedly biased, selection of degron substates proved amenable to rapid SspB-catalyzed degradation. We conclude that substrate-dependence of the SspB system remains a critical factor for the success of this degron system. For substrates that prove degradable, we provide a series of titratable SspB-expression vehicles.

## INTRODUCTION

Controllable inactivation of enzyme function has proven an invaluable tool in the functional study of essential gene products. The inhibition of essential protein function is usually affected by either specific chemical inhibitors binding to the enzyme active site (*1*), or by introduction of a genetic mutation conferring a conditional protein instability - usually to elevated temperatures (*2*). Using these and similar techniques, enzyme function may be controlled temporally in a way that is kinetically rapid, and occasionally reversible. Whereas essential genes may only be studied in vivo by this or similar methods, non-essential gene assays may also benefit from rapid gene product inactivation to uncover novel compensatory mechanisms, and where possible, genetic removal (knock-out) of a protein structural gene provides an ultimate reference point to conditional inactivation. Most cellular proteins possess a half-life orders of magnitude greater than the cellular generation time (*3*), such that gene knock-out produces loss of function by protein titration via cell doubling. Additionally, genetic tools, such as complementing a knock-out with a curable plasmid-borne copy of the gene, require additional cell doublings for plasmid titration before the protein concentration starts to fall. This slow loss of the targeted function allows time for potential cellular compensation mechanisms to ameliorate the original mutation’s effects, either by additional compensatory genetic mutations, or by metabolic adaptation, such that researchers may lose critical information concerning targeted gene function. Therefore, even for non-essential proteins, a strategy of rapid and controllable inactivation may provide additional insights unavailable to non-conditional approaches. Unfortunately, a generalizable, non-invasive, method for creating such specific protein deficiencies is lacking in many bacteria.

Generation of conditional alleles of a gene is time- and resource-intensive. Even in *E. coli*, the best studied and understood of all organisms, not more than perhaps 1% of its proteins may presently be studied by conditional inactivation. Other less well-studied organisms offer considerably fewer tools for conditional inactivation of gene functions. An alternative to the aforementioned approaches of chemical or genetic inactivation are the so-called “degron” systems of targeted protein destruction by cellular proteases (*4, 5*). In eukaryotes, a number of systems for targeted protein degradation have been developed with perhaps the predominant system leveraging a plant-specific auxin pathway for targeted degradation (*6*). Now in wide use in many eukaryotic organisms, including mammals, the system requires only the N- or C-terminal fusion of a short (∼70 AA) auxin-binding domain which upon binding auxin dimerizes with E3 ubiquitin ligase directing the ubiquitylation of the protein of interest, targeting it for degradation by the 26S proteasome. Similar systems have been reported to exist in Archaea (*7*), although to date none have been adapted for experimental use.

Bacteria, which lack a ubiquitin-like system to target proteins for destruction, are in general less amenable for directed protein degradation approaches. However, one system finding significant success in some bacteria is the tmRNA system for degron-tagging of incomplete proteins (*8–10*). Truncated mRNAs lack the stop codon necessary for ribosomal release, and so ribosomes are effectively trapped in unproductive complexes with truncated mRNA (*11*). To release and recycle stalled ribosomes, an alanine-charged tmRNA is recruited to the vacant A-site of the stalled ribosome. After the translocation of the tmRNA to the ribosome P site following addition of alanine to the truncated peptide, a template switch occurs, whereby the ribosome uses the tmRNA itself as mRNA to tag the truncated polypeptide and efficiently terminate translation. In *E. coli*, there is a single tmRNA gene named *ssrA*, which encodes 10 amino acid oligopeptide ANDENYALAA, terminated by two stop codons (*12*). The *ssrA* tmRNA appends 11 amino acids (the charged alanine, plus the tmRNA-encoded decapeptide) to the truncated protein and releases the trapped ribosome, while targeting the fusion protein for degradation by the ClpXP protease (*13*).

The ClpXP protease, one of the major *E. coli* protein-degradation complexes, fills many roles besides truncated protein destruction (*14*). Structurally, ClpP forms the core of the complex by assembling two stacked heptameric rings into a barrel shape with the protease active sites inside (*15*). Thus, the ClpP protease innards are not accessible to folded proteins or even short relatively linear peptides (*16*) and so ClpP protease requires active feeding. ClpX assembles into a hexameric planar ring containing an axial pore through which peptide chains are translocated in an ATP-dependent process (*15*). On its own, ClpX is characterized as an unfoldase or chaperone (*17*); however, when coupled to the ClpP protease, ClpX acts as a specificity factor by identifying and feeding unfolded substrate into the lumen of ClpP, where they are degraded (*18*). ClpX is able to exert at least ∼ 20 pN of force while unfolding a substrate, which should be enough to unfold most protein structures (*19*); however, the degree of force is influenced by the sequence of the peptide passing through the ClpX axial pore (*20*). It is possible for ClpXP to irreversibly disengage from a partially degraded substrate if the peptide within the ClpX channel exhibits poor “grip,” such as observed for poly-glycine, especially if this motif lies adjacent to a strongly folded domain (*21, 22*). In addition, the strength of ClpX binding itself adjacent to strongly folded domains influences its subsequent ability to unfold them (*23*).

The *ssrA*-encoded peptide contains two binding sites that direct and modulate its rate of degradation (*24*). The terminal LAA moiety binds directly to the ClpX unfoldase of the ClpXP protease, while AANDENY binds SspB, a substrate adaptor protein, which also binds ClpX, but to a different surface than the LAA tripeptide. In the context of the full-length tag, the presence of SspB serves to enhance delivery of SsrA-tagged substrates to ClpXP (*25*). Due to this redundancy in the SsrA tag, the absence of SspB results in no apparent phenotype in bacteria. The same cannot be said of ClpX-promoted degradation, responsible for complex cellular processes, like recovery from induction of the SOS-regulon (*26–28*), cellular division dynamics (*29*), and various stress responses (*30*). Therefore, appending a SspB-dependent version of the SsrA tag to proteins of interest should make their concentration regulatable by SspB availability in a physiologically non-invasive way.

Recognizing the potential importance of the SspB-specific binding determinant, the Sauer lab created *ssrA* tag variants, whose function would be completely dependent on SspB (*31*). Their altered *ssrA* tag, called “DAS+4”, retains the SspB binding domain, adds a 4 amino acid linker and changes the terminal LAA resides to DAS, allowing ingress of the peptide into ClpX but abolishing ClpX-specific binding. The ClpX / SspB / DAS+4 degron system has been deployed successfully in mycobacteria using a tetracycline-responsive promoter to drive expression of *E. coli* SspB (*9*). In *E. coli* a rapamycin-inducible degradation system employing a split-SspB has been reported (*32*), but displays a significant background degradation of the DAS+4-tagged target protein (*33*). To complement the split-SspB approach, we sought to develop an inducible degron system in *E. coli,* regulated by the expression of an intact SspB.

In this work, we present a general-purpose *ssrA*-based degron system in *E. coli* and characterize its major limitations. As SspB acts catalytically to deliver substrate to ClpXP, its expression must be tightly controlled to prevent premature target degradation. At the same time, a fast and robust SspB production is required to affect a near-complete degradation of the target protein upon induction. We engineered an SspB-variant with reduced stability and placed it under control of a strongly-inducible promoter – allowing for 1) no degradation in the “off” position and 2) a kinetically rapid “off” to “on” transition for SspB-mediated degradation of target proteins. We then employed this system to study a variety of enzymes involved in DNA metabolism. For “amenable” substrates, an apparently complete inactivation occurs within minutes of SspB induction. However, in many cases, depletion of enzymes involved in DNA metabolism was insufficient to result in a null phenotype. Surprisingly, in a few cases, the degron substrates appeared completely refractory to ClpXP degradation. With the help of substrates with easily quantifiable activities, like the classic reporters beta-galactosidase (beta-gal) or green fluorescent protein (GFP) (*34*), we found that native *E. coli* beta-gal (gp *lacZ*), while correctly recognized, appeared completely refractory to degradation by ClpXP. Another model degradation substrate, a fast-folding variant of GFP (*35*), was degradable by ClpXP, but if expressed from a strong promoter, failed to disappear completely in exponentially growing cells, highlighting the expected difficulty with robustly-expressed high-copy number proteins. We also identified a ClpX-independent mode of SspB-dependent protein inactivation. Finally, we found that SspB concentration modulates the rate of substrate degradation upon exit to stationary phase indicating a possibility that the DAS+4 tag maintains some ability to interact with the ClpAP protease.

## RESULTS

### Inducible SspB expression – The lactose promoter

To test how an inducible SspB-dependent degron system functions in *E. coli*, we first degron-tagged two well-understood DNA repair proteins, RecA and RuvB, with the “DAS#1” tag (Fig. 1A). Both RecA and RuvB are required for the full UV-resistance in *E. coli*, making UV survival a sensitive functional readout for our *recA* and *ruvB* constructs. At the same time, while RuvB has a modest copy number, RecA is a copious protein, — allowing us to gauge the degradation of both types of targets. We reasoned that tight control of SspB expression would be required, as the adaptor protein functions catalytically to deliver substrate to ClpXP (Fig. 1A). At the same time, we also desired a robust and kinetically rapid induction of SspB upon addition of inducer. The arabinose promoter is among the most tightly repressible of the inducible promoters (*36*), — however, its induction is kinetically slow (*37*). Therefore, we began investigations with a classical P*lac*-based expression vector, the pLAC22 plasmid, which contains the full-length lactose promoter and LacI repressor under control of the strongly upregulated *lacI*^q^ promoter (*38*). Together these elements provide sufficiently stringent conditions for *lac* promoter repression, while ensuring robust expression when inducer is present. However, even in the absence of inducer, pLAC22::*sspB* increased UV sensitivity of the *ruvB*-DAS#1 (Fig. 1B) and r*ecA*-DAS#1 (Fig. 1C) degron alleles beyond the empty vector, indicating significant SspB production.

**Figure 1.**
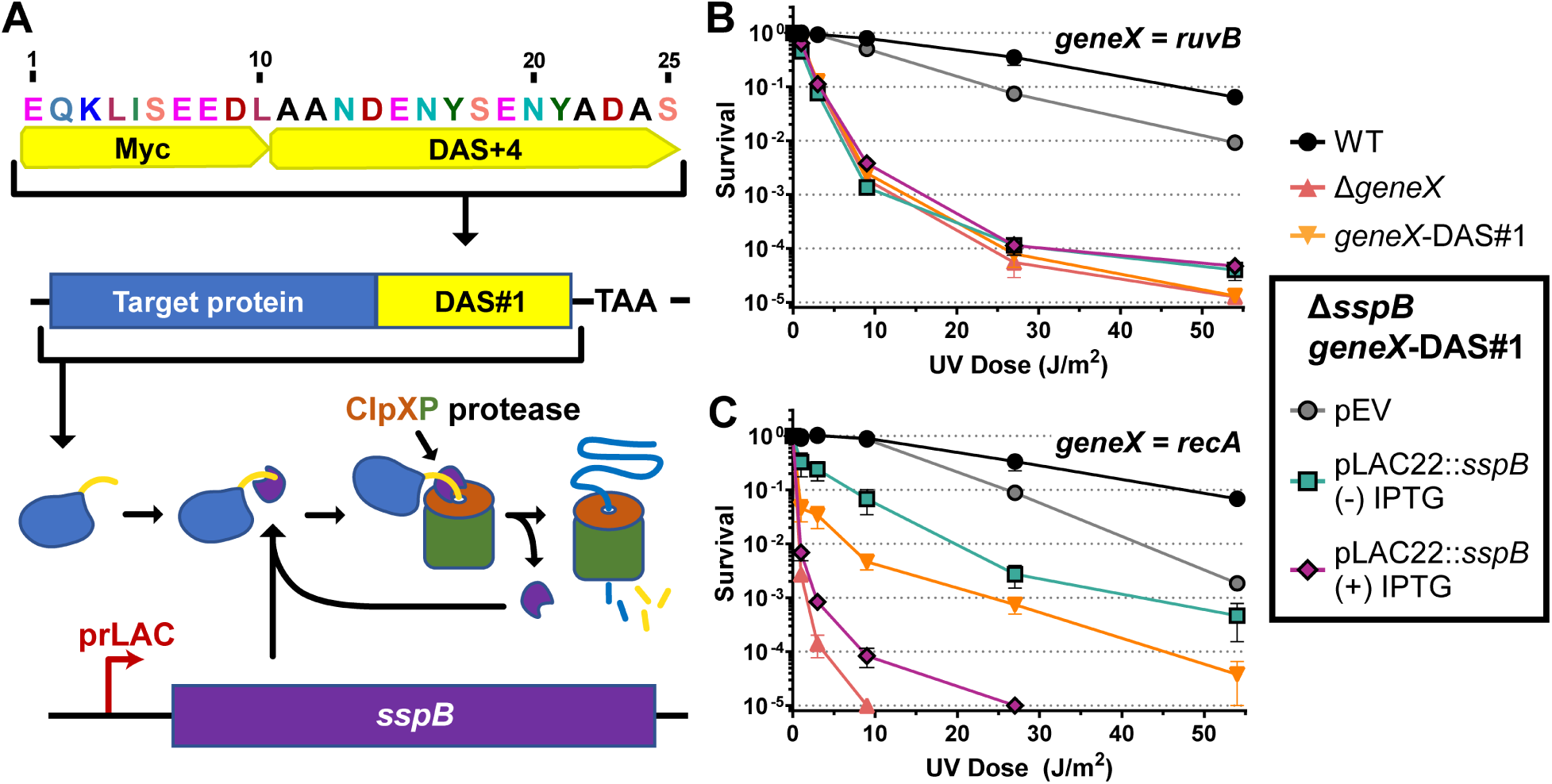
– A classic expression vector offers inadequate regulation of SspB production. **A.** Schematic of the “DAS#1” degron tag and mechanism of SspB-dependent degradation of the DAS+4 substrate mediated by ClpXP protease. **B.** UV sensitivity of the ΔsspB ruvB-DAS#1 strain carrying SspB on the pLAC22 plasmid containing the full-length lactose promoter and lacI^q^ repressor. **C.** UV sensitivity of the ΔsspB recA-DAS#1 strain carrying the same pLAC22::sspB plasmid from panel B. Plates contained 500 μg / mL ampicillin. Unless otherwise noted, here and throughout the paper, data are reported as the mean ± SEM of between 3 and 6 independent repetitions. Strains without pLAC22::sspB contain pLAC22 empty vector. WT is AB1157, Δ*recA* is JC10287, Δ*ruv* is AM547, *recA*-DAS#1 is eGC058, Δ*sspB recA*-DAS#1 is eGC059, *ruvB*-DAS#1 is eGC036, Δ*sspB ruvB*-DAS#1 is eGC061, pLAC22::*sspB* is pGC064.

In particular, in the case of the RuvB degron, the pLAC basal expression of SspB was sufficient to induce a fully-null *ruv* UV sensitivity (green squares) while the empty vector control (pEV, grey circles) displayed close to WT UV sensitivity, showing that the degron tag itself has almost no interference with RuvB function (Fig. 1B). The addition of inducer (purple diamonds) did not change this outcome; indicating that basal expression from pLAC22::*sspB* was already saturating for RuvB degradation. Often termed “leakiness”, low-level mRNA transcription from regulated promoters is a common phenomenon (*39*), and one which appears particularly problematic with catalytic proteins like SspB in combination with moderate protein copy number targets, like RuvB (200-400 molecules per cell) (*40*).

The high protein copy number target, RecA (3,000-10,000 molecules per cell) (*40*), behaved differently when tagged with DAS#1. The pLAC22::*sspB* driven expression showed a more nuanced situation, and the presence (purple diamonds) or absence (green squares) of inducer resulted in widely different UV sensitivities (Fig. 1C). Interestingly, induction of inducer produced a more severe UV sensitivity than seen for an empty vector strain producing SspB from the endogenous locus (orange inverted triangles), indicating that native levels of SspB are insufficient to affect a more complete turnover of high copy degron substrates. Additionally, this initial finding suggested that not only was tight repression of SspB expression required, but also a robust induction was necessary to achieve wild-type and mutant phenotypes in the uninduced and induced states, respectively.

### Inducible SspB expression – The tetracycline promoter

The tetracycline (TET) responsive promoter, P_LTETO-1_, seemed like a good choice for a SspB expression system as it offered a large dynamic range (*39*), and the added benefit of being responsive to the non-toxic tetracycline analogue, anhydrotetracycline (aTc)(*41*). To test SspB driven by P_LTETO-1_ we employed a slightly reconfigured degron tag consisting of a short linker, single myc-tag and SspB-specific DAS+4 peptide (Fig. 2A, “DAS#2”). When the *recA* and *ruvB* genes were fused to the corresponding sequence, we initially characterized them in the presence or absence of the chromosomally encoded *sspB*^+^. As was found for DAS#1, in the absence of the *sspB* gene, the *recA*-deg and *ruvB*-deg strains appeared wild-type, whereas *sspB*+ cells were either defective (*recA*, Fig. 2CD) or deficient (*ruvB*, Fig. 2EF).

**Figure 2.**
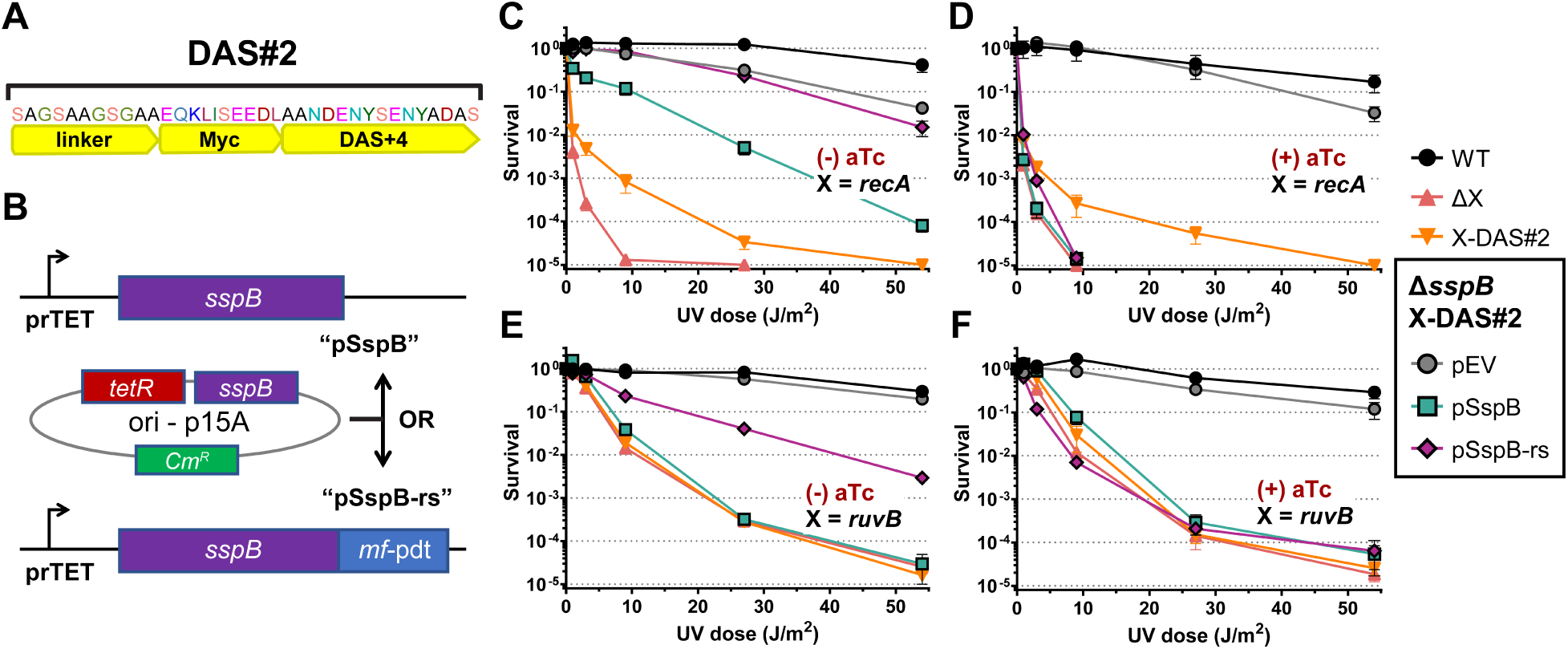
– A chimeric SspB possessing reduced basal expression. **A.** Schematic of the “DAS#2” degron tag. **B.** Cartoon schematic of the “pSspB” and “pSspB-<u>rs</u>” (reduced <u>s</u>tability) plasmids placing *sspB* under control of the tetracycline promoter. The plasmids are identical but for the fusion of a *Mesoplasma florum* protein degradation signal to the SspB C-terminus in Sspb-rs. **C-F.** UV-sensitivity for *recA*-degron (*recA*-nDAS) function (panels C and D) or *ruvB*-degron (*ruvB*-nDAS) function (panels E and F). The degron strains harbor empty vector (pEV), or *sspB* under control of a tetracycline-inducible promoter. Uninduced (panels C and E) lack addition of an inducer of SspB expression, while the non-toxic inducer anhydrotetracycline has been added in panels D and F. WT is AB1157, Δ*recA* is JC10287, Δ*ruv* is AM547, *recA*-DAS#2 is eGC566, Δ*sspB recA*-DAS#2 is eGC567, *ruvB*-DAS#2 is eGC568, Δ*sspB ruvB*-DAS#2 is eGC569, pEmpty is pZA31-luc, pSspB is pGC513-1, pSspB-rs is pGC533-1.

Interestingly, *recA*-DAS#2 displayed slightly increased sensitivity to chromosomal *sspB*, suggesting it was more efficiently degraded than *recA*-DAS#1. The p15A replicon of the plasmid-born *sspB* under control of the TET promoter, referred to as “pSspB”, has a reduced copy number relative to ColE1 replicon of pLAC22::SspB (25 vs. 60 copies per cell during exponential growth) (*39*). Despite the TET promoter’s increased dynamic range, with pSspB, a decreased gene function (increased UV sensitivity) was still observed without inducer. Leaky expression from P_LTETO-1_-SspB was more apparent when paired with *ruvB*-DAS#2; we observed a complete loss of Ruv function in this background similar to that seen with *ruvB-*DAS#1 pLAC22::SspB (Fig. 2EF, “pSspB”, green squares). Accounting for the higher efficiency of DAS#2 degradation, we may infer a somewhat decreased basal expression for pSspB compared to pLAC22::*sspB*.

### Tighter regulation of SspB by a stability-reducing tag

One common strategy to decrease basal steady-state expression is to amend the protein of interest with a degron-tag in order to lower its half-life. Initially, we fused the full-length SsrA or ClpX-specific LAA recognition tags to SspB but found by semi-quantitative assays that SspB-SsrA lacked any SspB function, and SspB-LAA while competent to deplete RuvB was unable to robustly deplete RecA (Fig. S1). Apparently, such sub-optimal SspB function resulted from the rapid degradation of the SspB-SsrA and SspB-LAA proteins by ClpXP. As an alternative to a ClpXP-specific degron, the heterologous SsrA-like protein tag from *Mesoplasma florum* (*42*) offered an alternative protein-destabilization element. Fused to GFP, the *M. flourm* SsrA-like tag (AANKNEENTNEVPTFMLNAGQANYAFA, “pdt”) directed a mild turnover of GFP-pdt by endogenous *E. coli lon* protease (*43*). In preliminary experiments with the Mf-lon system we verified this “background” degradation of a *pdt* tagged GFP absent its cognate (MF-lon) protease (*43*). We then fused *pdt* to the carboxy-terminus of SspB in pSspB, making pSspB-rs (reduced <u>s</u>tability) and found that, as desired, pSspB-rs displayed a significantly reduced leakiness without induction (Fig. 2C and E, compare pSspB to pSspB-rs), such that *recA*-deg was now fully resistant to UV treatment, and *ruvB*-deg, while still showing some UV sensitivity, was now significantly improved compared to the Δ*ruvB* mutant. At the same time, pSspB-rs still provided a robust and rapid induction of SspB when aTc was added (pSspB-rs, compare Fig. 2C vs D and E vs F). Upon observing this success from the plasmid-based system, we integrated the expression elements from SspB-rs into the genome in place of the native *sspB* gene. While our integrated expression construct was identical in composition and structure to pSspB-rs, lacking only the plasmid replication origin, it failed to produce a sufficient repression of P_LTETO-1_ (Fig S10).

### A system of SspB expression vectors for differentially expressed targets

To further repress uninduced SspB levels in the ruvB-deg background, we decreased the strength of the ribosomal binding site (RBS) of pSspB-rs sspB mRNA. Using web-based RBS prediction software, we systematically produced a series of plasmids containing incrementally weaker predicted RBSs (*44*). As can be seen in Figure 3, pSspB-rs-rbs2 showed WT levels of UV sensitivity even with *ruvB*-deg allele, while still allowing for sufficient expression to achieve the null phenotype in the induced state. Since pSspB-rs already showed the required performance with *recA*-DAS#2, we did not test different RBS plasmids with it.

**Figure 3.**
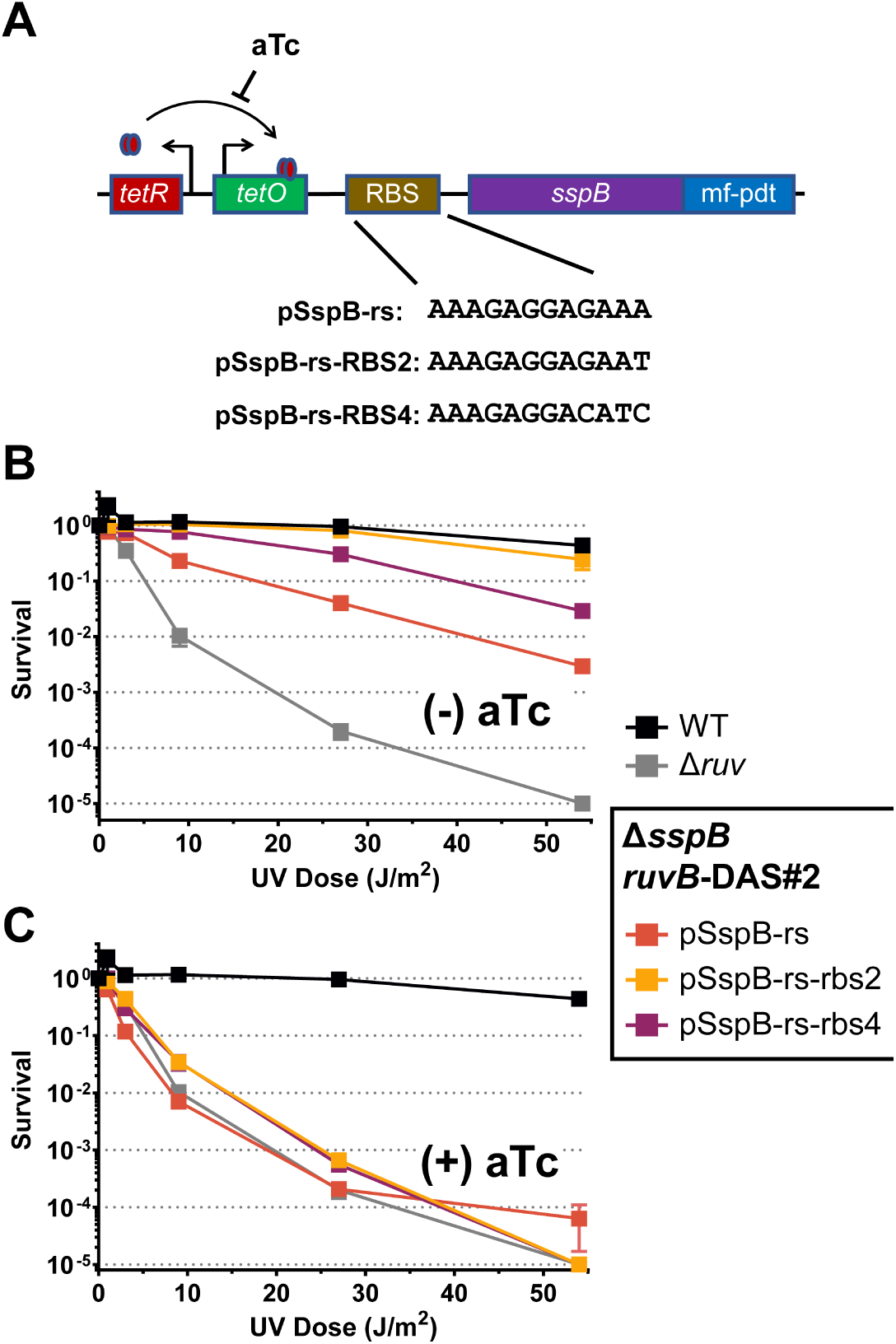
– Ribosomal binding site mutations further reduce basal expression from pSspB-rs. **A.** Schematic of the pSspB-rs promoter region and nucleotide sequences for the ribosomal binding sites (RBS) of the unaltered and “RBS-down” pSspB-rs derivatives shown in panels B and C. **B.** UV-sensitivity assay of *ruv*-degron harboring the parental pSspB-rs or the derivative pSspB-rs-rbs2 or -rbs4 plasmids without addition of inducer. **C.** UV-sensitivity assay of the same strains as in B, but with aTc inducer added. WT is AB1157, Δ*ruv* is AM547, Δ*sspB ruvB*-DAS#2 is eGC569, pEmpty is pZA31-luc, pSspB-rs is pGC533-1, pSspB-rs-rbs2 is pGC589, pSspB-rs-rbs4 is pGC591.

### SspB-dependent target inactivation may be ClpX-independent

We tested ClpX-dependence of the *recA*-deg and *ruvB*-deg phenotypes at a single UV dose of 30 J (m^2^)^-1^ such that *recA*-DAS#2 and *ruvB*-DAS#2 in the presence of genomic levels of SspB should appear null for gene function. As expected, the *ruvB*-DAS#2 strain became UV-sensitive upon SspB expression in a *clpX*^+^ strain (Fig. 4A, blue bars), while its *clpX^-^* sibling was entirely UV-resistant even when SspB was highly expressed (Fig. 4A, red bars).

**Figure 4.**
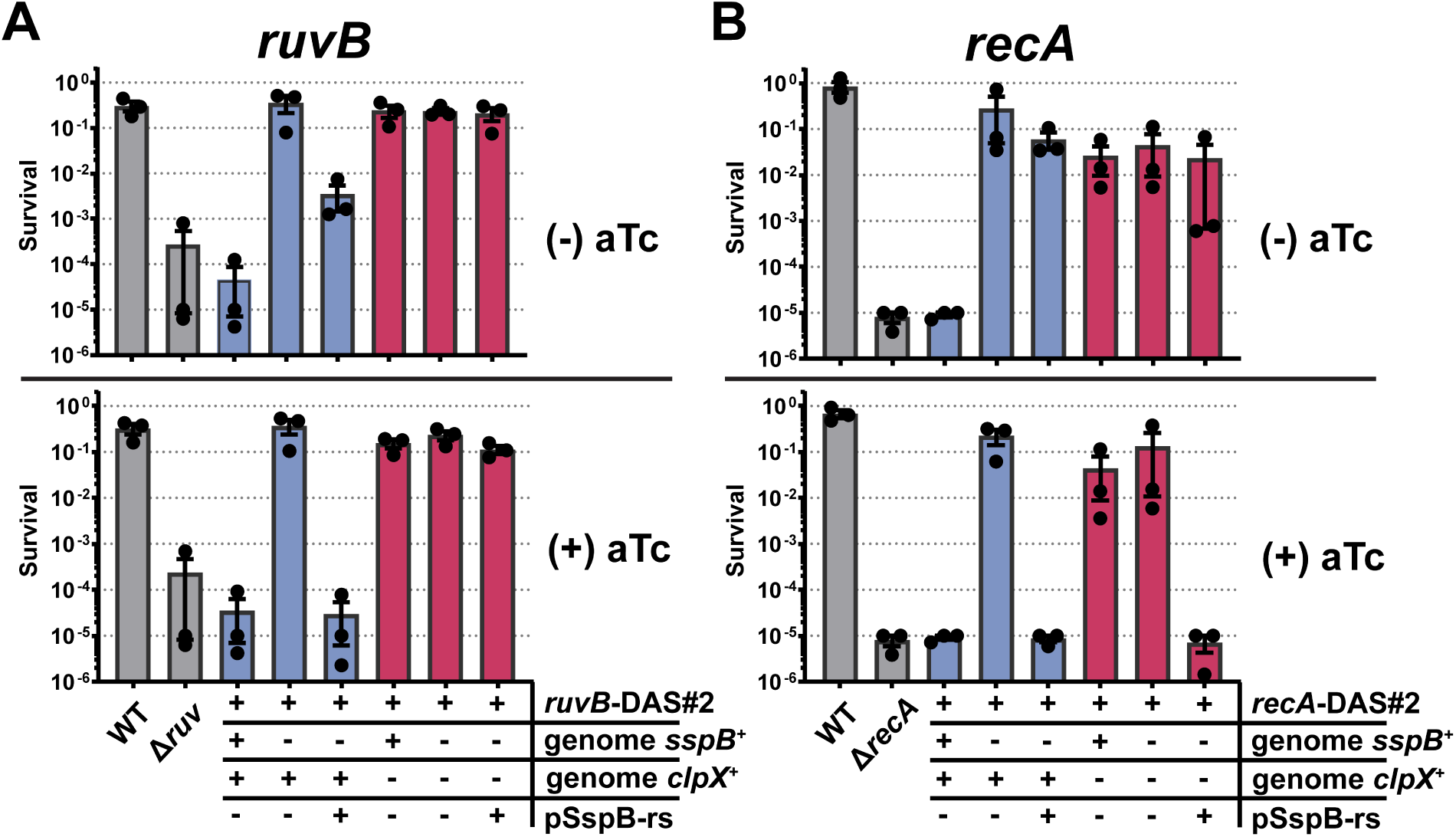
– Inactivation of RecA-degron is partially independent of the ClpX AAA+ ATPase. **A.** UV-sensitivity assay with exposure to 30 J/m2 of UV, normalized untreated controls. “Genomic sspB” indicates the status of the endogenous genomic copy of *sspB*, and “clpX” the endogenous genomic copy of *clpX*. **B.** An identically constructed experiment as in A, but for *recA*-degron. All strains lacking pSspB-rs contain empty vector. WT is AB1157, Δ*recA* is JC10287, Δ*ruv* is AM547, *recA*-DAS#2 is eGC566, Δ*sspB recA*-DAS#2 is eGC567, Δ*clpX recA*-DAS#2 is eGC682, Δ*clpX* Δ*sspB recA*-DAS#2 is eGC683, *ruvB*-DAS#2 is eGC568, Δ*sspB ruvB*-DAS#2 is eGC569, Δ*clpX ruvB*-DAS#2 is eGC684, Δ*clpX* Δ*sspB ruvB*-DAS#2 is eGC685, pEmpty is pZA31-luc, pSspB-rs is pGC533-1.

In contrast to RuvB-DAS#2, even in the Δ*clpX* background, *recA*-DAS#2 could still be inactivated by SspB – but, only when SspB was expressed at high levels. In Figure 4B, a differential response to SspB expression levels can be seen between Clp+ and Clp– backgrounds. While a genomic copy of sspB is sufficient to render ClpX+ cells *recA* deficient, in a *clpX*-*sspB*+ background the same *recA*-DAS#2 allele appears fully functional (Fig. 4B, compare the leftmost blue and red bars). At the same time, induction of SspB expression from pSspB-rs completely removes RecA-DAS#2 function in both ClpX+ and ClpX– backgrounds (Fig. 4B, bottom panel, compare the rightmost blue and red bars). Loss or alteration of function of the DAS#2 tagged RecA or RuvB was ruled out by complementation of *clpX*-with a plasmid-borne copy of ClpP/X (Fig. S2). The likely explanation for ClpX-independent RecA-deg inactivation is that SspB binding to RecA-DAS#2 protein interferes with polymerization of RecA necessary for its function, — however due to the differential requirements for SspB ± ClpX, this pathway cannot be the only one, and likely operates alongside the ClpX-dependent (degradation) mechanism of inactivation, providing another way to inactivate some high copy number target proteins, dependent on SspB overproduction.

### Degron tagging of multiple DNA replication and repair proteins

To test the general applicability of our apparently robust degron/SspB system, we built a number of degron alleles of DNA replication and repair genes and assessed the resulting mutant phenotypes. Surprisingly, completely functional “gold-standard” alleles similar *recA*-deg and *ruvB*-deg formed a minority of our, admittedly biased, list of targets. As can be seen in Table 1, of the 15 genes tested, two lost function when C-terminally tagged even in the absence of SspB. Five of the fifteen, or 30% of all candidates, showed no phenotype upon SspB induction, although all displayed a correct DAS+4 fusion when sequenced. An additional four of the fifteen displayed partial phenotypes of varying degree upon SspB induction. This left only three fully functional fusions, *recA*-deg, *ruvB*-deg and *dnaG*-deg, which behaved as WT in the uninduced state and at the same time became fully mutant upon SspB induction. With so many degron alleles failing to respond to SspB induction, we sought a better understanding of what could be preventing SspB-mediated degradation.

**Table 1.** – Degron alleles tested in this study. Protein amount are for Neidhardt EZ rich defined medium from (*40*), +++, complete inactivation; ++, 50% inactivation; + measurable effect.

| Gene | Copy Number | Tags tested | Functional? | Inactivated? | Tested by |
| --- | --- | --- | --- | --- | --- |
| RecA | 10377 | DAS#1, #2, #3 | + | +++ | UV sensitivity |
| RuvB | 452 | DAS#1, #2 | + | +++ | UV sensitivity |
|  |  | DAS#3 | + | ++ | UV sensitivity |
| dnaG | 356 | DAS#2 | + | +++ | Viability |
|  |  | DAS#3(no myc) | - | N/A | N/A |
| PolA | 2028 | DAS#2 | + | + | UV sensitivity |
|  |  | DAS#3(no myc) | + | ++ | UV sensitivity |
| RecB | 93 | DAS#2 | + | - | UV sensitivity |
|  |  | DAS#3(no myc) | + | ++ | UV sensitivity |
| GFP | N/A | DAS#2, #3 | + | ++ | Flourescence, Western |
| PriA | 105 | DAS#2 | + | + | UV sensitivity |
|  |  | DAS#3(no myc) | - | N/A | N/A |
| RecC | 118 | DAS#2, #3(no myc) | + | - | UV sensitivity |
| RnhA | 804 | DAS#3(no myc) | + | - | UV sensitivity |
| HolD | 338 | DAS#3 | + | - | AZT sensitivity |
| lacZ | 31 | DAS#2, #3(no myc) | + | - | B-gal assay, Western |
| LigA | 2004 | DAS#2, #3(no myc) | + | - | UV sensitivity |
| dnaA | 840 | DAS#2, #3(no myc) | - | N/A | N/A |
| HolC | 266 | DAS#2 | - | N/A | N/A |

### Beta galactosidase is completely resistant to SsrA-dependent degradation

How a particular mutant phenotype reflects a corresponding protein’s abundance is not well understood for the vast majority of proteins, with an obvious exception of those few that have to polymerize in long filaments, like RecA. With this in mind, we chose to employ degron substrates that provided a quantitative readout of protein abundance. Two such proteins, that work as monomers, are beta-galactosidase (beta-gal, LacZ) and green fluorescence protein (GFP), which can be assayed by colorimetric assay (*45*) or by fluorescence (*46*), respectively. While GFP degron fusions have been explored by a number of other researchers as ClpXP-degradable substrates, to our knowledge, no a beta-gal-deg allele has been reported in the literature.

For our work with these quantitative proteins, we started with DAS#2 tag, but collected quantitative data with a third DAS+4-based degron tag, DAS#3 (Fig. 5A). The DAS#3 tag is considerably longer than DAS#1 and DAS#2 tags and contains a long alpha-helical linker region to spatially separate the substrate from the myc-DAS+4 tail. The linker region allows for degradation of the c-myc tag by ClpXP even if lacZ cannot be degraded, as 28 residues remain adjacent to domains resistant to denaturation by ClpXP (*47–49*). Additionally, during the course of these studies, important determinants of ClpX unfoldase function were found to reside in the substrate’s peptide sequence (*20*). As ClpXP is engaged with a substrate, there are five amino acid residues residing within the ClpX pore, termed a “grip” sequence which strongly enhances the rate at which ClpX unfolds stably folded domains. Therefore, SspB-dependent GFP degradation was tested for three different linkers of sufficient length with different reported secondary structures, two rigid and one flexible (*50*), in combination with an efficient grip sequence (“acidic” grip sequence, (*20*)), and found a rigid alpha-helical linker to provide the best expression and good SspB-dependent degradation characteristics (Fig. S3). Additionally, we compared the efficiency of DAS#3 to DAS#2 with respect to RecA- and RuvB-degron UV sensitivity and found that DAS#3 functioned adequately in those contexts, although with slightly reduced efficiency compared to DAS#2 (Fig. S4).

**Figure 5.**
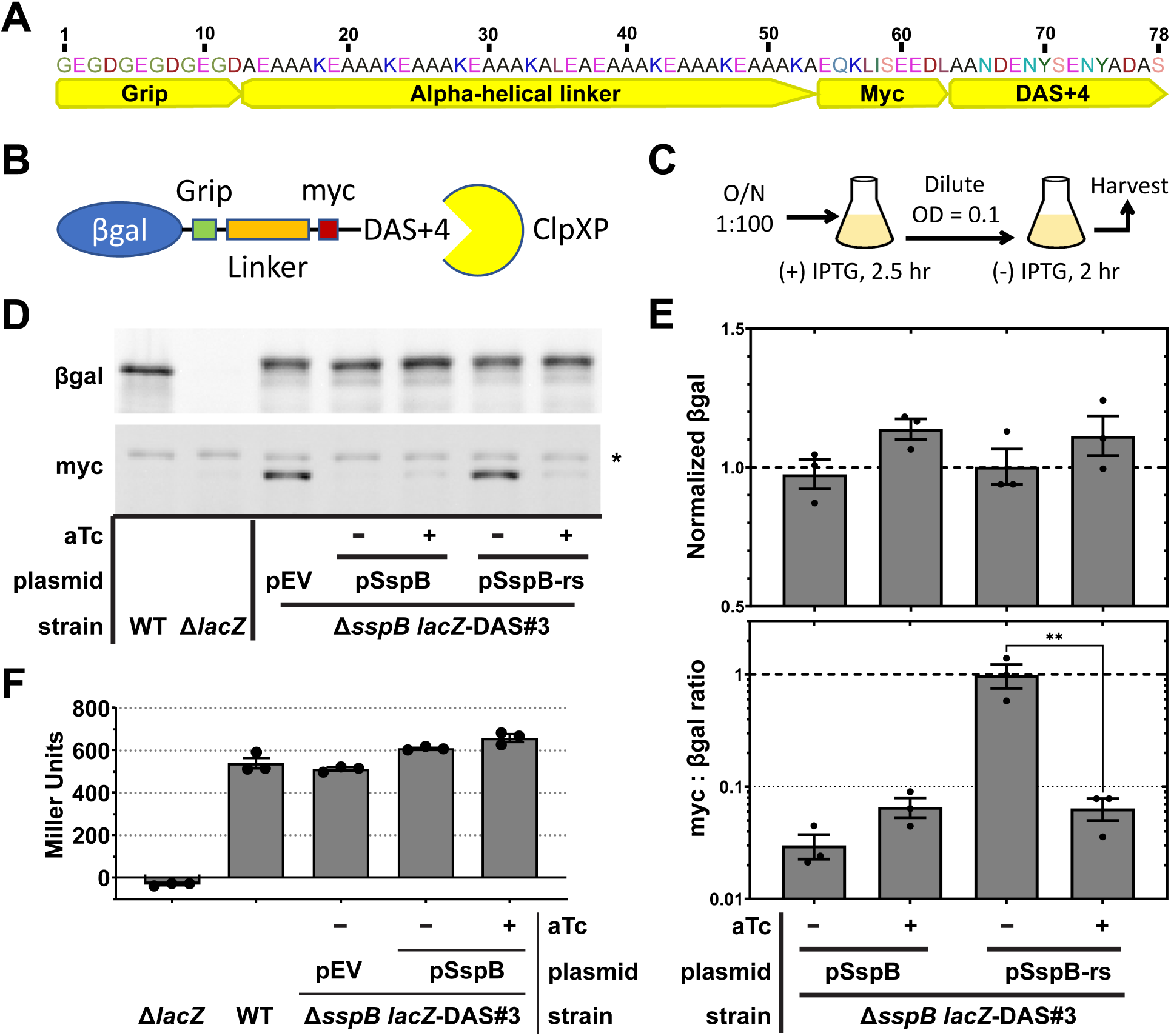
– Beta-galactosidase is refractory to SspB-mediated degradation. A. Schematic representation of the “DAS#3” degradation tag. The “grip” sequence has been reported to increase the unfolding ability of ClpX, — see the text for details. The alpha-helical linker was intended to increase the ability of ClpXP to degrade the DAS+4 and myc moieties. **B.** A cartoon representation of lacZ fused to the DAS#3 tag with ClpXP depicted degrading DAS+4. **C.** Experimental overview depicting the “pulse” of beta-gal expression, allowing time for its degradation absent significant new synthesis. **D.** A representative western blot, one of the three, used to generate the aggregate data shown in panel E. **E.** Upper panel, western blot beta-gal signal normalized to the WT signal (Panel D, lane 1). Lower panel, ratio of the myc : beta-gal western blot signals, i.e. the “nibble” ratio. The myc signal was normalized to empty vector (Panel D, lane 3), before calculating the ratio. Note that myc signal alone is not reported as it appears almost identical to the ratio panel (The ratio panel results from dividing by ∼1). WT is MG1655 pZA31-luc, Δ*lacZ* is JT33, Δ*sspB lacZ*-DAS#3 is eGC706, pEV is pZA31-luc, pSspB is pGC513-1, pSspB-rs is pGC533-1.

Appending DAS#3 to LacZ did not alter either beta-gal amount (Fig. 5D) or activity (Fig. 5E, the upper panel). Even the induced amount of SspB did not affect the specific beta-gal activity (Miller Units, Fig. 5F), making one wonder if beta-gal-DAS#3 was at all recognized as a substrate by ClpXP. Using replicate western blots, we probed for LacZ or the Myc epitope of the DAS#3 tag, while varying the amount of intracellular SspB. The disappearance of Myc confirmed that ClpX initiated processing of beta-gal-deg in a SspB-dependent manner, but the LacZ structure was still completely refractory to degradation. When normalizing band intensity to a strain completely lacking SspB (pEV = empty vector), leaky expression from pSspB, or induced expression of SspB from pSspB or pSspB-rs was sufficient to reduce the Myc signal by ∼20-30 fold (Fig. 5DE). Only the uninduced pSspB-rs yielded a Myc : beta-gal ratio near unity. At the same time, the beta-gal signal by western blot, like the beta-gal enzymatic assay, showed no significant difference for any condition (Fig. 5D). Thus, some proteins are resistant to ClpXP degradation, naturally limiting the usefulness of this degron approach.

### Degradation of GFP does not keep pace with its production

To measure the kinetic parameters of the DAS#3 degron tag, we employed a fast-folding GFP variant, GFPmut3b (*35*)(henceforth referred to as GFP), which is frequently used in degron studies (*43, 51–55*). Repression of *lacZ* expression is mediated by the LacI repressor binding to three *lacO* operator sites, one of which, O_2_, lies within the *lacZ* coding sequence (*56*). To mirror as closely as possible the LacZ-deg result in Figure 5, we replaced the *lacZ* coding sequence with that of *gfp* (Fig. 6A). To our surprise, we observed this normally tightly-controlled promoter to be constitutively active, such that GFP protein levels failed to react appreciably to addition of the inducer IPTG (Fig. S5). Moreover, during log-phase growth, GFP levels were reduced by only ∼20% when SspB was induced from pSspB-rs (Figure 6E, compare T=0 for grey, blue, and red bars), suggesting that the rate of GFP (replenishing) synthesis was close to the rate of SspB-catalyzed GFP degradation.

**Figure 6.**
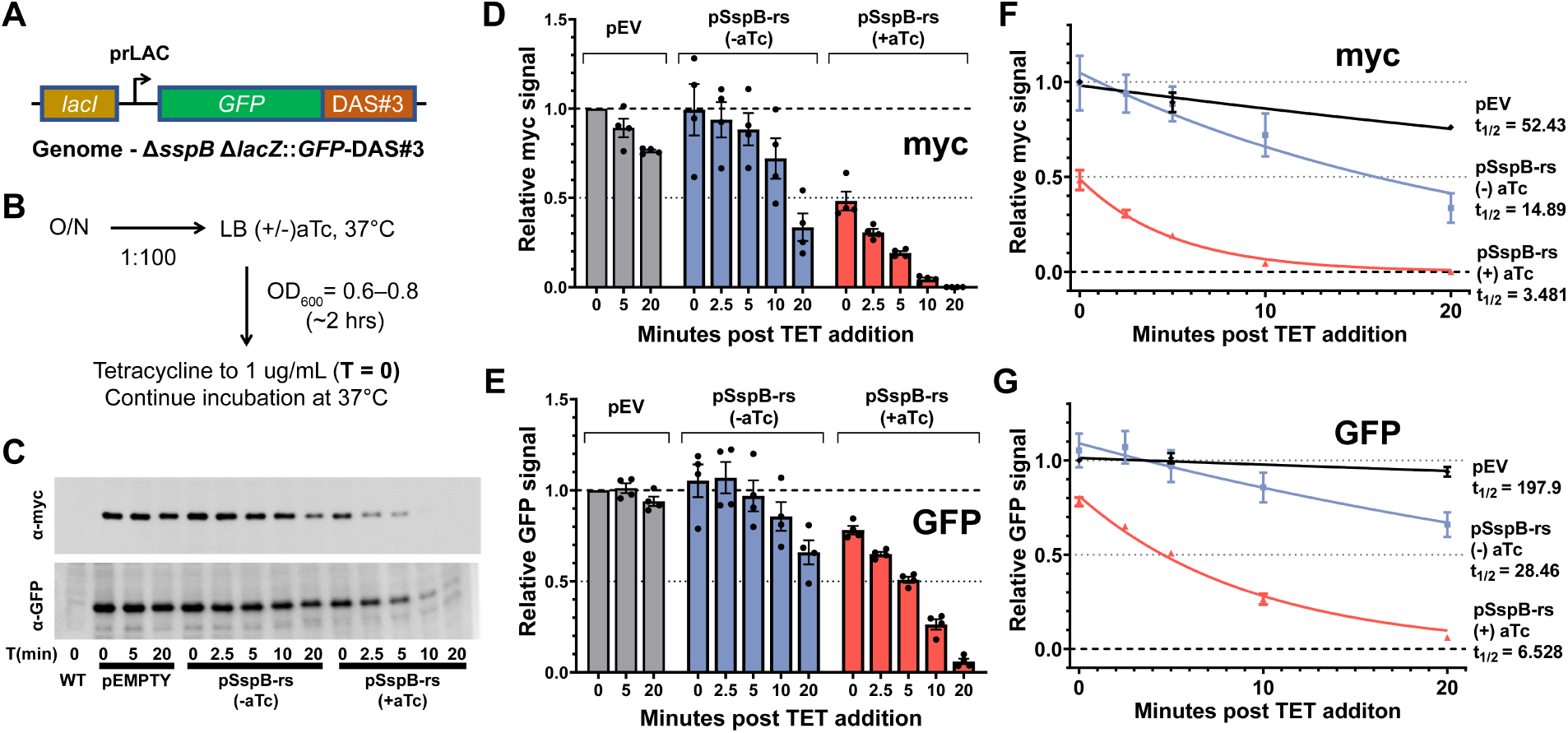
– Kinetics of GFP-degron degradation. **A.** Schematic of the modified *lacZ* locus : replacement of LacZ coding sequence with that of GFP. **B.** Experimental overview of GFP pulse by termination of protein synthesis by addition of tetracycline (since GFP expression at the *lacZ* locus is constitutive). **C.** A representative western blot; several such blots were used to generate the summary data shown in panels D-G. **D.** Western blot myc signal at different timepoints following tetracycline addition. **E.** Western blot GFP signal at the same timepoints following tetracycline addition as in panel D. **F.** Kinetic half-life measurement curves and values for western blot myc signal. **G.** Same as in F but for GFP. WT is MG1655 pZA31-luc, Δ*sspB* Δ*lacZ:*:GFP-DAS#3 is eGC762, pEmpty is pZA31-luc, pSspB is pGC513-1, pSspB-rs is pGC533-1.

To avoid this complication of the analysis of degron function by new protein synthesis, rather than minimizing synthesis by withdrawal of IPTG inducer, as was done for beta-gal, we blocked new protein synthesis by addition of tetracycline. As can be seen in Figure 6 D-G, contrary to the degradation resistance of beta-gal, GFP was readily degraded in a SspB-dependent manner. Fitting the data with a one-phase decay indicated a myc signal half-life of 3.5 minutes, while GFP signal dropping by half in 6.5 minutes (Figure 6, FG). We note that our 1x myc tag is recognized by a monoclonal antibody, while the GFP antibody was polyclonal, so these small differences in half-life likely reflect the time required to either unfold or completely degrade GFP, and our results are broadly coincident with other recently reported GFP-degron half-life measurements (*57*). We also note that uninduced pSspB-rs affected a lower, but still measurable turnover of myc and GFP – a point that is better addressed in the following section.

Similarly to the LacZ functional assay, we confirmed our western blot findings with a GFP functional assay. Since our detection methods were not sensitive enough for direct detection of GFP fluorescence when GFP expression was driven from the chromosomal *lacZ* promoter (Fig. S6), we followed the lead of others and drove GFP expression from the constitutive *lacI*^q^ promoter housed on a ColE1 replicon-containing plasmid (*43*). The “pGFP-DAS#3” plasmid expressed ∼16-fold more GFP than the genomic *lacZ*::*gfp* construct (Fig. S7). In this context, GFP signal was sufficient to allow fluorescence detection in live cell culture during continuous growth in a plate reader. These more detailed kinetic data, like the western result, confirmed the SspB-dependent degradation of GFP (Fig. 7). Interestingly, the GFP half-life measurements displayed a more than 2-fold increase compared to the western data in Figure 6, suggesting that ClpXP capacity may be limiting when GFP-DAS#3 is expressed from this plasmid expression system. Nevertheless, the GFP fluorescence and western blot data are in good agreement and indicate that GFP-DAS#3 is an amenable degron substrate, while DAS#3 is a functional degron tag — confirming beta-gal as a refractory substrate.

**Figure 7.**
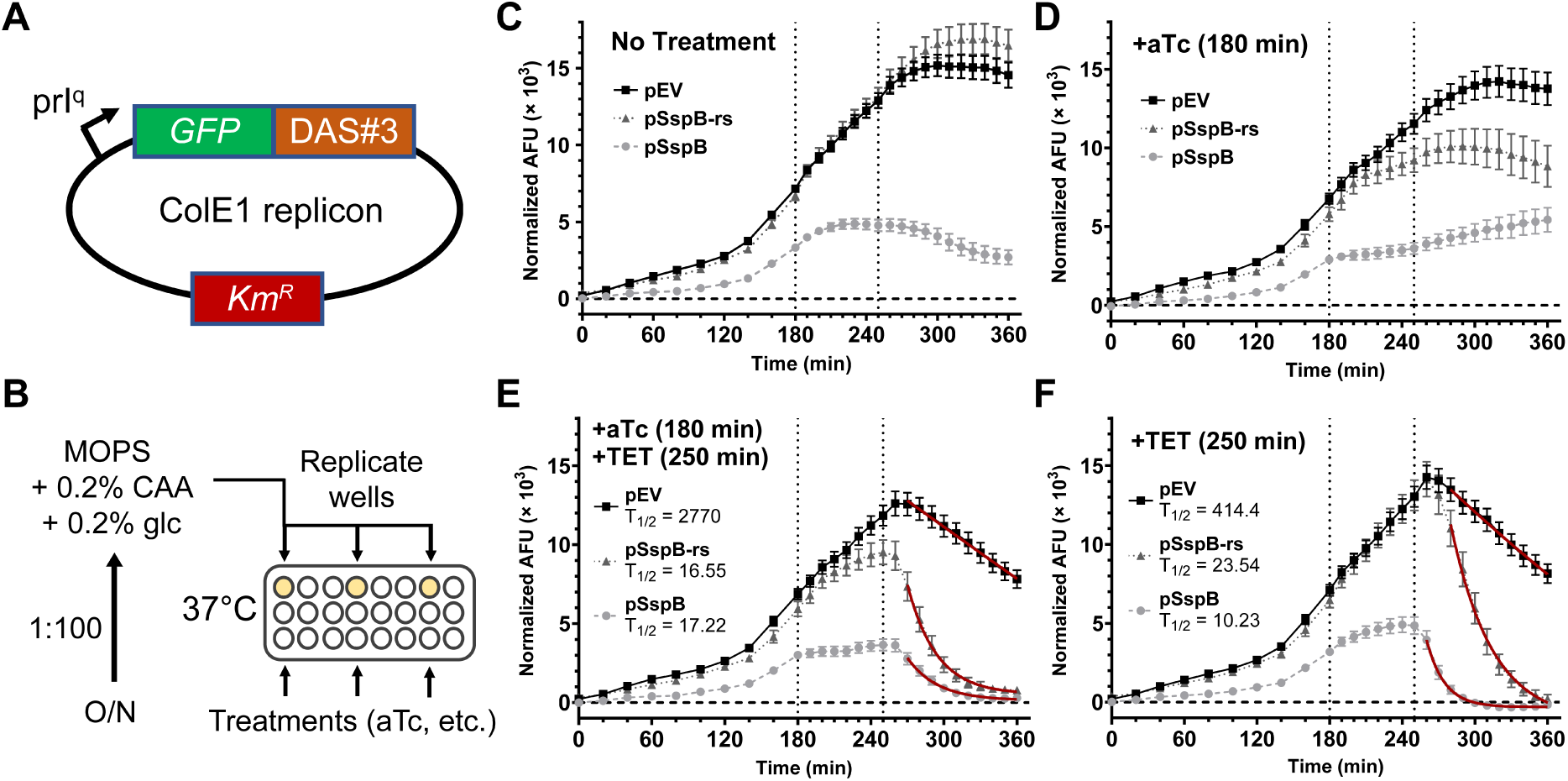
– Kinetic analysis of GFP-DAS#3 degradation in live cells. **A.** Schematic of the GFP-expression plasmid yielding ∼50-fold higher GFP expression than the chromosomal expression shown in Figure 7. **B.** Schematic of experimental design depicting the replicate cultures used as controls or subjected to different treatments. **C-F.** Background subtracted (autofluorescence) mean fluorescence measurements for empty vector, pSspB and pSspB-rs strains containing the constitutively expressing GFP plasmid.

Vertical dotted lines indicate the times of addition of anhydrotetracycline (aTc, the inducer of SspB, left line) or tetracycline (TET, a protein synthesis inhibitor, right line), although both treatments were added only in panel E. Panel C had no treatment, while panel E cells were treated only with aTc, while panel F, TET only. Data points represent means of 12-24 independent measurements ±SEM. Red lines indicate the fitted one-phase decay function yielding the half-life values given in each panel. Growth curves and doubling times for all conditions are provided in Fig. S8. The WT strain used for autofluorescence normalization was eGC706 pZA31-luc. Strain eGC027 hosted the pGFP-DAS#3 plasmid (pGC707-1) in addition to one of: pEV (pZA31-luc), pSspB (pGC513-1) or pSspB-rs (pGC533-1).

### Excess SspB interferes with target protein degradation

The SspB adaptor protein physically tethers the degron substrate to the ClpXP protease. Initially we had assumed that substrate-ClpX binding (and therefore the rate of degradation) would become saturated at some concentration of SspB relative to substate, such that further increases in SspB concentration would no longer affect the kinetics of substrate degradation. Unexpectedly (but perhaps related to the “Hook Effect” (*58*)), the *gfp*-DAS#3 allele, in addition to degradation with intermediate SspB levels (Fig. 7, pSspB-rs), revealed that too much SspB can inhibit substrate degradation. In Figure 8 we present evidence in support of a “local maximum” for SspB-dependent degradation, using the original *sspB* allele (not the reduced stability variant). The pSspB expression vector is significantly leaky even when uninduced, — so that it provides enough SspB to direct the degradation of RuvB (Fig. 2) or the LacZ-DAS#3 myc tag (Fig. 6). We therefore found it odd when we observed a small but consistent increase in substrate concentration (RuvB, UV sensitivity) upon induction of pSspB (Fig. 2, compare pSspB in panels E and F) (Fig. 5, panel D compare lanes 4 and 5, and panel E compare pSspB ± aTc).

**Figure 8.**
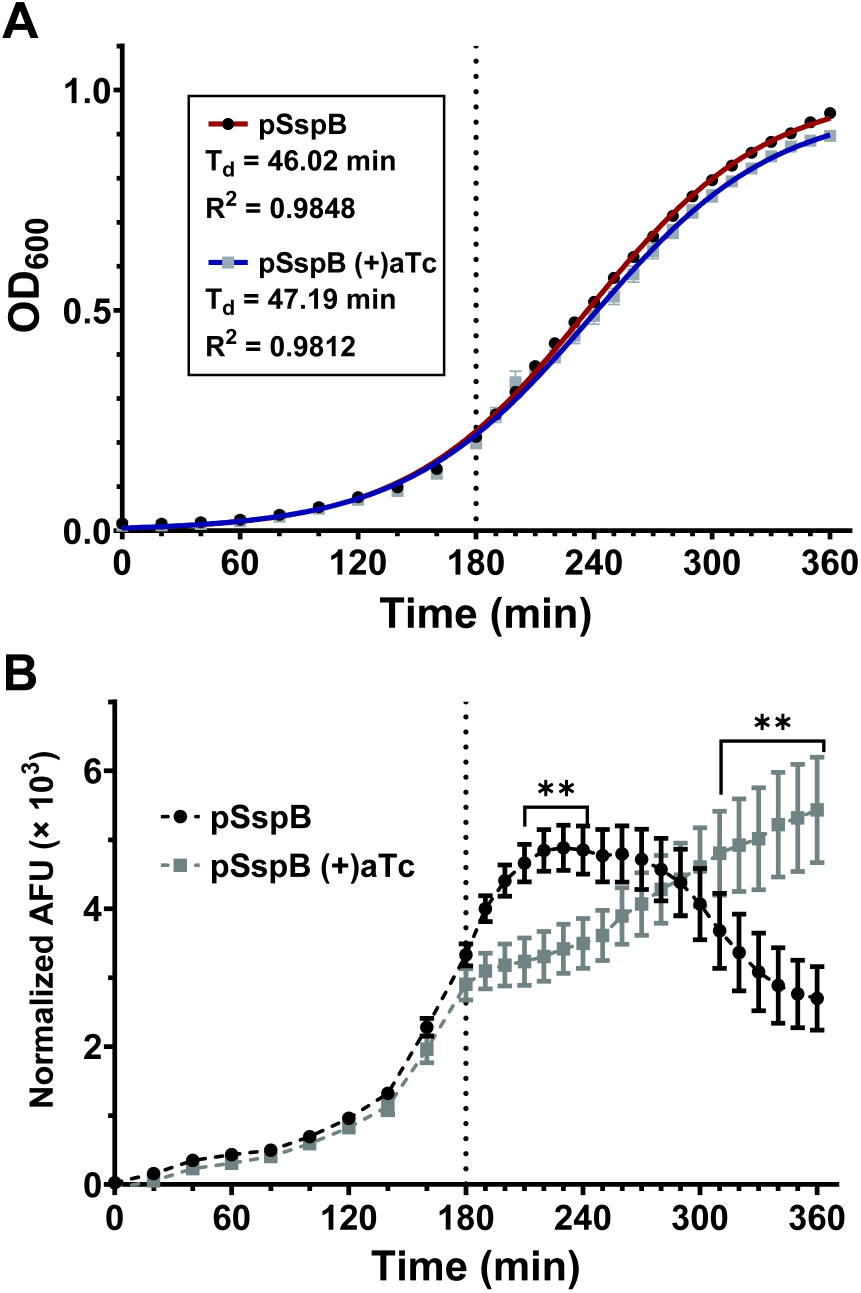
– Increased SspB expression negatively affects substrate levels during entry to stationary phase. **A.** OD600 measurement of cell growth for the strain and conditions shown in panel B, with the best fit logistic growth curves in red and blue and the resulting fit values and doubling times given in the inset. **B.** The evolution of GFP fluorescence following addition (grey squares) or not (black circles) of anhydrotetracycline at T = 180 min., reproduced from data in Figure 7, panels D and C. Asterixis represent P > 0.01 for paired t-tests. Error bars are SEM with n = 12 (+)aTc, or n=24 (-)aTc. The vertical dotted line highlights the time of addition of anhydrotetracycline (aTc).

The more granular live cell GFP data produced statistically significant demonstration of this effect. Figure 8 contains duplications of two fluorescence curves from Figure 7 (Panel B) and adds the growth curves for these strains in panel A. As can be seen, aTc is non-toxic at the concentration used throughout this work, as aTc did not significantly affect growth rate (only the 360 min. difference is significant, also see Figs. S8 and S9). While growth rate is unaffected, clear differences in GFP expression can be seen between cultures uninduced (black circles) or induced (grey squares) for (the unmodified) SspB expression. While the expected reduction in GFP is observed following aTc addition, these relative differences disappear and then invert at later timepoints, indicating that high levels of SspB may be inhibitory to the degradation of SsrA-tagged substrates, especially as cells approach stationary phase. It has been reported that the SsrA tag is degraded by the ClpAP protease as well as ClpXP, and that ClpAP is induced upon entry into stationary phase (*55*). Additionally, binding of SspB to the wild-type SsrA tag is known to inhibit SsrA degradation by ClpAP (*24*). Therefore, it seems reasonable to postulate that our DAS#3 tag is degraded by ClpAP, although several of the ClpA-interacting residues are missing in the DAS+4 tag.

## DISCUSSION

The degron system for controlled protein elimination is a great asset for any organism, in which it works reliably. The goal of this work was to create such a general-purpose and broadly-applicable SsrA-based degron system in *E. coli*. We leveraged a protein destabilization element from a very different microbe, *Mesoplasma florum*, to destabilize the uninduced SspB adaptor protein to minimize its basal expression. We then drove SspB expression from the robust P_LTETO-1_ promoter, creating a kinetically rapid SsrA-based degron system. Despite the inclusion of the SspB-destabilization element, low-abundance degron target proteins remained at least partially depleted in the presence of the uninduced SspB plasmid. Therefore, we created several additional plasmid variants containing weakened RBS elements to better titrate background SspB expression, while allowing full off-to-on depletion of poorly-expressed substrates.

Employing plasmids from this series we demonstrated precise control of the RecA and RuvB DNA repair proteins, which both showed complete stability absent inducer - and at the same were completely inactivated within minutes upon induction of SspB synthesis. However, when we sought to extend these results to other DNA repair enzymes, or quantitative model substrates, we observed widely variable effects: some substrates proved manipulatable, while others were resistant to SspB-catalyzed degradation. Thus, a general ClpXP-based degron system may not be practical in *E. coli*, because a significant fraction of its proteins show surprising stability against degradation.

Two proteins with quantitative phenotypes, GFP and beta-gal, revealed challenges to a generalized SsrA-based system in *E. coli* for controlled protein degradation. A fast-folding variant of GFP is degradable both *in vitro* and *in vivo* by ClpXP (*52, 55*). However, during exponential growth in a rich medium at 37°C, GFP-DAS#3 produced from the genomic *lacZ* locus was not degraded quickly enough in the presence of SspB to reduce its steady-state expression by more than two-fold. Oddly, our *lacZ*::GFP construct, under control of the native *lacZ* promoter, was relatively unresponsive to the *lac* operon inducer IPTG and showed less than 2-fold induction in the presence of saturating concentrations of IPTG (Fig. S5). This sharply contrasts with the 300-fold induction reported for beta-gal induced under similar conditions We suspect that the loss of the O_2_ operator site within the *lacZ* coding sequence is at least partially responsible for the observed dysregulation (*56*). If GFP expression in our uninduced cells approximates the number of beta-gal monomers observed in induced cells (*59*), then the number of GFP molecules may be assumed to be ∼3-6 x 10^3^ per cell. A copy number of 3000-6000 per cell is common in bacterial physiology (*40*). Endogenous ClpXP was reported to possess the capacity to degrade this quantity of GFP, and also keep up with ongoing synthesis: *In vivo*, using the wild-type SsrA sequence, a plasmid-borne GFP-SsrA expressed at a copy number of 100,000 per cell was degraded to completion (*55*). In our system, significant GFP-DAS#3 remains in the presence of SspB + ClpXP, and the T_1/2_ values for GFP and the DAS#3 c-myc tag are quite similar (Fig. 7), suggesting that SspB-mediated recruitment of DAS#3 to ClpX is rate-limiting. Replacement of the rigid alanine linker with a rigid proline or flexible linker did not change this result (Fig. S3) (*50*).

A GFP functional assay using a plasmid-borne copy of GFP-DAS#3 buttressed the western blot results. As GFP fluorescence may be assayed in live cells, this assay allowed for a more granular examination of GFP degradation throughout the growth phase. Again, as seen with the western results, induction of SspB expression from pSspB-rs reduced GFP fluorescence by less than two-fold (Fig. 7D), and a block to protein synthesis (Fig. 7 E, F) was required for GFP to be degraded to the detection limit. Examination of GFP levels in strains containing the initial pSspB plasmid, lacking the SspB-destabilization element, produced a puzzling result. In Figure 8 it can be seen that induction of pSspB, as expected, reduces GFP levels. However, this effect is transient and by late log / early stationary phase the induced strain possessed significantly more GFP than the uninduced culture. This concentration-dependent inhibitory effect for SspB-catalyzed degradation has previously been observed in the context of the wild-type SsrA tag, where SspB inhibited GFP-SsrA degradation by the ClpAP protease (*24*). The ClpAP binding residues defined by Flynn et al (*24*), are only partially present in the DAS+4 tag. Additionally, Farrell et. al (*55*) observed a hand-over of protease activity to ClpAP in stationary phase, however ClpAP levels did not increase until a cell density of OD_600_ = 2-3 was reached; far beyond the inhibitory threshold of OD_600_ = 0.75 seen in our experiments. Despite these differences, we consider it possible that a ClpXP independent degradation signal is present in DAS+4, and is masked by SspB binding. An alternative explanation is that SspB expression from the pSspB plasmid increases in late log phase and that this excess free SspB protein competitively inhibits binding of substrate-loaded SspB, as observed *in vitro* Why this effect was not observed with RuvB-degron is unknown, but its existence for at least some substrates presents a problem for a generalized system, as many proteins, if reduced by only 50% like GFP, would accomplish most or all of their functions, — and therefore the corresponding cells would still appear WT.

If a substrate is too numerous or difficult to degrade, ClpXP complexes may become saturated (*61*). Therefore, the “degradability” of a ClpXP substrate may also prove critical to degron system function. One quantitative substrate tested, beta-gal, encoded by the *lacZ* gene, proved this point, as beta-gal appeared completely refractory to degradation by ClpXP. Substrate engagement by ClpXP was demonstrated by the quick, SspB-dependent, loss of the DAS+4-proximal Myc tag; at the same time, no loss of beta-gal activity or antigenic signal was observed (Fig. 5). Cases of non-degradable substrates have been described in the literature, but these usually have a glycine rich motif adjacent to a well-folded domain (*20*). When a poly-glycine containing peptide resides in the ClpX axial pore, the peptide-translocating loops have difficulty exerting sufficient force to unfold the adjacent domain, and occasionally translocates backwards far enough to disengage from the substrate. Beta-galactosidase shows no obvious change in mobility by western analysis; however, we considered the possibility that a glycine motif may exist close to the C-terminus and allow for escape from ClpXP by a known mechanism. The most C-terminal residue of beta-gal, lysine 1024, begins a beta-strand integral to the core structure of domain 5 (*62*). Disruption of this terminal beta strand occurs in *lacZ* nonsense mutant X90 (*63*) where ∼12 amino acids are truncated from the end of the beta-gal protein (*64*) resulting in its rapid degradation and a null phenotype (*65*). This degradation of the truncated form is particularly striking, as it has been demonstrated that beta-gal is normally so stable as to be only diluted by cell division with no apparent proteolysis (*66*). Since no glycine residues are found in the terminal 11 residues, it would appear highly unlikely that a glycine-slippage mechanism protects beta-gal from degradation by ClpXP. That the X90 mutant protein is unstable suggests that LacZ cannot be unfolded by ClpXP at all and displays complete resistance despite being provided with a well characterized strong “grip” sequence. To the best of our knowledge, only one other (non-glycine mediated) substrate refractory to degradation by ClpXP is the *E. coli* dihydrofolate reductase protein (*49*).

In addition to poorly degraded or non-degraded degron substrates, we also identified a ClpX-independent effect of SspB binding to the DAS+4 tag. This effect was seen for the RecA protein, which functions as a polymer, and we consider it likely that SspB binding blocks RecA polymerization. To function in recombinational repair and SOS induction, RecA must polymerize on damaged DNA to form a nucleoprotein filament capable not only of guiding a homology search, but also of cleaving the LexA transcription repressor (*67*). Cleavage of LexA induces the expression of the “SOS response,” comprising the coordinated expression of a number of DNA repair proteins including RecA itself (*67*). Thus, RecA polymerization is a critical component of the cell’s high-capacity DNA damage repair system.

The degron system we describe was designed as a generalized tool for targeted protein degradation, but fails to degrade most targets to phenotype-assessed completion. Putting aside proteins which for one reason or another lose function when C-terminally tagged, many degron-tagged targets appear to maintain a significant concentration due to their relatively high rates of synthesis and/or resistance to proteolysis by ClpXP. The interplay between these rates of synthesis and degradation, in the presence of SspB, dictates the target’s steady-state concentration and consequent phenotype. Unfortunately for the system described above, many tested substrates managed to avoid a phenotypically-complete degradation. However, for amenable substrates this tool provides a unique genetic approach to conditional protein depletion, which is both kinetically similar to, and experimentally compatible with, classical temperature-sensitive gene alleles, and thus provides a useful tool for genetic analysis.

## MATERIALS AND METHODS

### Strains media and growth conditions

Two wild-type *E. coli* K12 strains were used in this work, MG1655 and a close relative, AB1157 – genotype and references are in the strain list. AB1157 is the preferred background for studies of DNA damage – first being used by the Howard-Flanders group due to its apparent genome stability when challenged with severe DNA-damaging treatments. The AB1157 genome was recently sequenced (NCBI RefSeq accession, NZ_CP076404.1). Strains were cultured in LB (Lysogeny Broth) containing, per liter, 10 g Tryptone, 5 g Yeast extract, 5 g NaCl and adjusted to pH 7.2 with 250 µl of 4 M NaOH. Solid medium was LB with 15 g of agar added per liter. Where indicated, strains were cultured in the presence of the following concentrations of antibiotics: 100 μg/mL ampicillin (Ap), 5 μg/mL kanamycin (Km), 10 μg/mL tetracycline (Tc), 10 μg/mL chloramphenicol (Cm). Unless otherwise noted, standard concentrations of antibiotics were present in all media used for propagation of plasmid-bearing strains. Where noted, the inducers anhydrotetracycline (aTc, Cayman Chemical #10009542) and isopropyl β-D-1-thiogalactopyranoside (IPTG, Research Products International #I56000) were added at 100 ng/mL and 1 mM, respectively. Data for the titration of aTc induction is provided in Fig. S9.

### Strain and plasmid construction

Design of strain and plasmid constructions is given in the supplemental primers list. Where primers are provided for genome editing, the method of Datsenko and Wanner was employed (*68*). Otherwise, the method of cloning is provided alongside the primer sequences.

### UV survival assay

Overnight cultures were diluted 1:100 and grown with shaking at 28 °C to early-/ mid-exponential phase (OD_600_ = 0.2-0.4). Serial dilutions of cultures in 1 % NaCl were spotted by 10 µl in replicate on solid media. Immediately after spots had dried (∼ 5 minutes), plates were irradiated in a Hoefer UVC 500 UV Crosslinker which had been masked to reduce UV flux to 1 J (m^2^ · s)^-1^ as measured by a UVC Digital Light Meter (model UV512C, General). All manipulations were performed under light from a yellow lamp (F15T8-GO, General Electric) to avoid photoreactivation. Irradiated plates were incubated in the dark at 28 °C for 14 – 16 hours before the pin-prick colonies of survivors were counted under a dissection microscope. Survival curves were constructed by normalization to the titer of unirradiated control within each experiment before calculation of mean and standard error between experiments.

### Beta-galactosidase assay

Saturated overnight cultures grown at 28 °C without IPTG were diluted 1:100 into LB + Cm + IPTG and shaken at 37 °C to OD_600_ = 0.8 – 1.0 (∼ 2.5 hours). After measuring OD_600_, one milliliter of each culture was washed two times (using aspiration to remove the supernatant) with an equal volume of 1% NaCl and diluted to OD_600_ = 0.1 in 5 mL of fresh media lacking IPTG. The OD_600_ of the diluted cultures were measured, and 1 mL was withdrawn, cells pelleted by centrifugation at 16,000 x *g* for 3 minutes and pellets frozen at -80 °C. The remaining 4 mL of each culture was returned to 37 °C and incubated again to OD_600_ = 0.8 – 1.0 (∼ 2 hours). After measuring OD_600_, 0.5 mL of each culture was pelleted as before and frozen at -80 °C. Frozen pellets were thawed and resuspended in 100 μL of 1% NaCl and added to 2 ml of Z-buffer (100 mM NaPO_4_ pH 7.0, 10 mM KCl, 1 mM MgSO_4_, 50 mM β-mercaptoethanol (βME),

0.00125% SDS) in a glass tube. Twenty-five microliters of chloroform was added, mixed by vigorous vortexing, and cells allowed to lyse while shaking for 10 minutes at 28°C. 240 μg of *ortho*-Nitrophenyl-β-galactoside (ONPG) was added (From a freshly made stock of 4 mg/mL in Z-buffer), and incubation continued at 28 °C for 10 minutes. Reactions were terminated by addition of 1 mL of 1 M Na_2_CO_3_, and Beta-galactosidase activity (Miller Units) was calculated as described (*45*).

### Beta-galactosidase western blot

After growth in liquid culture as detailed in the “Beta-galactosidase assay” section above, cells equating to 1.5 OD_600_ units were pelleted and frozen at -80 °C. Cell pellets were thawed, resuspended in 100 μL SDS-PAGE buffer (80 mM Tris pH 6.8, 2% SDS, 10 % glycerol, 0.0006% BPB, 2.5% βME), boiled for 10 minutes, centrifuged at 16,000 x *g* for 10 minutes and 40 μL of supernatant removed to a fresh tube and mixed by pipetting. Fifteen-microliter portions of each sample were electrophoresed on freshly poured duplicate polyacrylamide gels. Gels were comprised of a 4% acrylamide, pH 6.8 stacking layer and 10 % acrylamide, pH 8.8 separating layer, both containing 0.1 % SDS and made from acrylamide / Bis-acrylamide solution (37.5:1, Biorad) according to established methods (*69*). Electrophoresis was performed at 50 V for 10 minutes followed by 200 V for 35 minutes. Gels were then electroblotted at 0 °C in Towbin buffer (*70*) to PVDF membrane (Immun-Blot, Biorad) for 600 minutes at 30 V. The quality of protein transfer was visualized by staining with Ponceau S (0.1% dye in 5% acetic acid, Sigma) and then blocked with 5% nonfat instant milk (Carnation) in PBS (137 mM NaCl, 12 mM Phosphate, 2.7 mM KCl, pH 7.4) for 1 hour at room temperature. Primary antibody was applied in PBST (PBS + 0.1% Tween 20) at room temperature for 1 – 1.5 hours. Blots were then rinsed with PBST and washed 3 times for 10 minutes each with PBST before adding secondary antibody in PBST and incubating at room temperature for one hour. Blots were rinsed and then washed 3 times for 15 minutes in PBST, after which the horseradish peroxidase (HRP) signal was visualized by applying “atto” substrate (SuperSignal West Atto, ThermoFisher) with imaging on an iBright CL1000 (ThermoFisher).

Of the two replicate gels, one was probed with a 1:3,000 dilution of α-betagalactosidase rabbit polyclonal antibody (ThermoFisher, A-11132), and secondary a 1:200,000 dilution of goat α-rabbit HRP (Abcam, ab205718). The second replicate gel was probed with a 1:3,000 dilution of α-myc mouse monoclonal antibody (Abcam, ab32) and secondary 1:200,000 dilution of goat α-mouse HRP (Abcam, ab97023).

### GFP western blot

Saturated overnight cultures grown at 28 °C were diluted 1:100 into 20 mL of LB + Cm with or without aTc and shaken at 37 °C to OD_600_ = 0.6 – 0.8 (∼ 2 hours). Once cultures reached the target OD_600_, the T = 0 timepoint was taken, Tc added, and the cultured returned to 37 °C. For each timepoint, to stop protein synthesis and degradation, 2 mL of the culture was added to 2 mL of LB, 20 mM KCN, 10% pyridine (*71*) at 0 °C, and the treated cultures were kept on ice until all timepoint were collected. Cultures were pelleted, and the pellets frozen at -80 °C. Pellets were then processed as for the Beta-galactosidase western blots except that 10 μl of lysate was loaded per well. Antibody dilutions, blotting and imaging were also identical to the beta-galactosidase experiment, except that rabbit polyclonal α-GFP (ThermoFisher 50430-2-AP) was used in place of α-betagalactosidase.

### GFP fluorescence assay

Strains were grown overnight at 28 °C to saturation in MOPS minimal medium (*72*), supplemented with 1.12 mM K_2_HPO_4_, 0.2% glucose, 0.1% casamino acids, 50 μg / mL thiamine, with Cm and Km. The next day cultures were diluted 1:100 into the same medium, and 300 μL added to wells of a black-walled, clear-bottomed 96-well plate (ThermoFisher M33089) for incubation in a Cytation 1 multi-mode plate reader (Agilent BioTek). Plate lids were treated with 10% Triton X-100 in ethanol to prevent fogging (*73*). Incubation in the plate reader was at 37 °C using continuous double-orbital shaking at the most vigorous setting. GFP signal was collected using a gain of 45 and read height of 10.25 mm. GFP and OD_600_ were read every 20 minutes for 3 hours, at which time aTc was added from a 50x stock in 25% EtOH. The plate was returned to incubation, and readings collected every 10 minutes for 1 hour. Tc was added from a 50x stock in 25% EtOH, and datapoints collected every 10 minutes for 1 hour, which marked the end of the experiment. We found it necessary to perform anhydrotetracycline and tetracycline additions sequentially (a single draw and multiple dispenses) from a multichannel electronic pipet (Eppendorf) while leaving the 96-well plate in the holder of the reader. Although this method caused sample carry-over between wells, we considered such effects were likely negligible and indeed none were ever observed. The alternative method of sequential dispenses, with tip changes to prevent carry-over, required the plate to sit for several minutes outside the machine causing substantial lags in both growth and GFP-expression kinetics (not shown). In total, sequential injection of aTc or Tc took no longer than 45 seconds. Preliminary experiments showed plate “edge” effects to be significant and 1% NaCl, or media blanks, were placed in columns 1 and 12. For three independent experiments, four biological replicates of each of the four strains were arrayed into five groups of two columns each. The “no addition” control was represented twice, as technical replicates, in both columns 2 + 3 and 10 + 11. The treatment groups: aTc, aTc / Tc, and Tc resided in columns 4 + 5, 6 + 7 and 8 + 9 respectively, such that addition of the aTc or Tc occurred in adjacent columns.

Preliminary experiments showed MOPS media, unlike LB, possessed low intrinsic fluorescence. However, even when grown in MOPS, *E. coli* displayed a strong, density-dependent, autofluorescence in the GFP channel (Fig. S6). Therefore, after subtracting media-specific blanks, the GFP autofluorescence signal observed for the WT control (lacking GFP) was used to normalize the signal from experimental strains by subtraction of the per-timepoint GFP signal (not by interpolation as used in Fig. S6). Additionally, the normalization was performed only within treatment groups (no treatment, aTc, aTc / Tc, Tc), to account for treatment-specific growth rate differences, and only within independent experiments. OD_600_ data were adjusted by subtraction of the media blank. Following normalization, the data were combined to generate summary statistics.

### Statistical Analyses

Densitometric band intensities were calculated in ImageJ (version 1.53e) using the “Plot Lanes” tool and suggested best practices (https://imagej.nih.gov/nih-image/manual/tech.html). Kinetic half-lives were calculated in GraphPad Prim (version 9.5.1) using the “one-phase decay” function with default settings. The doubling times and model curves reported in Figure 8 were calculated in GraphPad using the “logistic growth” function with default settings. Significant differences were also calculated in GraphPad: For Figure 5, a ratio paired t-test was employed with default settings, and for Figure 8 multiple paired t-tests were employed, without correction for multiple comparisons, and otherwise default settings.

## AUTHOR INFORMATION

### Corresponding Author

**Glen E. Cronan –** University of Illinois at Urbana-Champaign, Department of Microbiology, University of Illinois at Urbana-Champaign, Urbana, IL 61801, USA

### Author

**Andrei Kuzminov –** University of Illinois at Urbana-Champaign, Department of Microbiology, University of Illinois at Urbana-Champaign, Urbana, IL 61801, USA

## AUTHOR CONTRIBUTION

A.K. conceived the project. A.K. and G.E.C. designed the experiments and wrote the manuscript. G.E.C. performed the experiments and analyzed the data.

## SUPPORTING INFORMATION

**Figure S1:** SsrA and LAA tagging of SspB; Figure S2: Complementation of *clpX* deletions; Figure S3: Alternative degron tag linkers; Figure S4: DAS#3 tagging of *recA* and *ruvB*. Figure S5: Constitutive expression of Δ*lacZ::GFP*: Figure S6: Negligeable fluorescence from Δ*lacZ::GFP*-DAS#3; Figure S7: Comparison of plasmid- and chromosomal-encoded GFP constructs; Figure S8: Growth curve and doubling times for experiments of Figure 7; Figure S9: Titration of induction by anhydrotetracycline; Figure S10: Comparison of basal SspB expression from pSspB-rs and Δ*sspB*::SspB-rs; Table S1: Strains; Table S2: Plasmids; Table S3: Primers and construction details (PDF).

## ACKNOWLEDGMENTS

We thank the entire Kuzminov lab for helpful discussions and feedback, Elena Kuzminova for creation and testing of the *rnhA* degron allele, and John E. Cronan for the *lacZ* deletion strain.

## For Table of Contents Use Only

Degron-controlled protein degradation in Escherichia coli: New Approaches and Parameters Glen E. Cronan* and Andrei Kuzminov

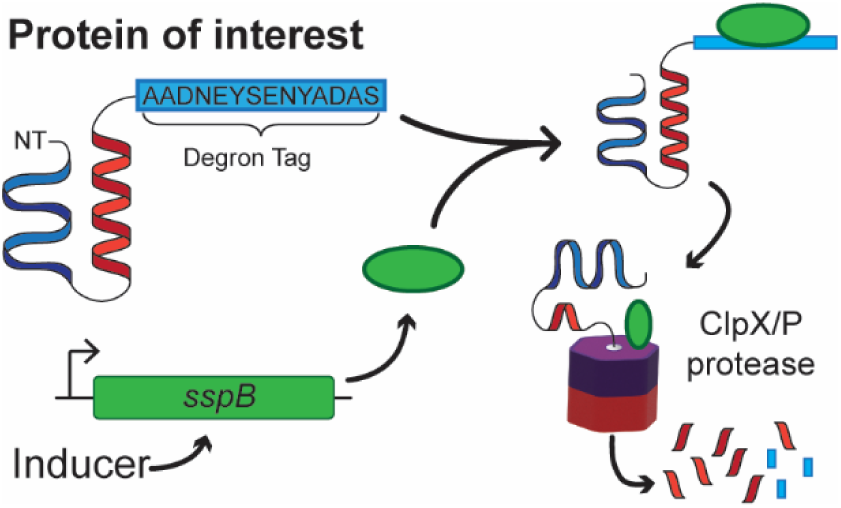

## REFERENCES

1. Knight, Z. A., and Shokat, K. M. (2007) Chemical genetics: where genetics and pharmacology meet, Cell 128, 425–430.

2. Pringle, J. R. (1975) Induction, selection, and experimental uses of temperature-sensitive and other conditional mutants of yeast, Methods Cell Biol 12, 233–272.

3. Koch, A. L., and Levy, H. R. (1955) Protein turnover in growing cultures of Escherichia coli, J Biol Chem 217, 947–957.

4. Izert, M. A., Klimecka, M. M., and Górna, M. W. (2021) Applications of Bacterial Degrons and Degraders - Toward Targeted Protein Degradation in Bacteria, Front Mol Biosci 8, 669762.

5. Natsume, T., and Kanemaki, M. T. (2017) Conditional Degrons for Controlling Protein Expression at the Protein Level, Annu Rev Genet 51, 83–102.

6. Nishimura, K., Fukagawa, T., Takisawa, H., Kakimoto, T., and Kanemaki, M. (2009) An auxin-based degron system for the rapid depletion of proteins in nonplant cells, Nat Methods 6, 917–922.

7. Fu, X., Liu, R., Sanchez, I., Silva-Sanchez, C., Hepowit, N. L., Cao, S., Chen, S., and Maupin-Furlow, J. (2016) Ubiquitin-Like Proteasome System Represents a Eukaryotic-Like Pathway for Targeted Proteolysis in Archaea, mBio 7.

8. 8. Griffith, K. L., and Grossman, A. D. (2008) Inducible protein degradation in Bacillus subtilisusing heterologous peptide tags and adaptor proteins to target substrates to the protease ClpXP, Molecular Microbiology.

9. Kim, J. H., Wei, J. R., Wallach, J. B., Robbins, R. S., Rubin, E. J., and Schnappinger, D. (2011) Protein inactivation in mycobacteria by controlled proteolysis and its application to deplete the beta subunit of RNA polymerase, Nucleic Acids Res 39, 2210–2220.

10. Badrinarayanan, A., Lesterlin, C., Reyes-Lamothe, R., and Sherratt, D. (2012) The Escherichia coli SMC complex, MukBEF, shapes nucleoid organization independently of DNA replication, J Bacteriol 194, 4669-4676.

11. Keiler, K. C., and Feaga, H. A. (2014) Resolving nonstop translation complexes is a matter of life or death, J Bacteriol 196, 2123–2130.

12. Tu, G. F., Reid, G. E., Zhang, J. G., Moritz, R. L., and Simpson, R. J. (1995) C-terminal extension of truncated recombinant proteins in Escherichia coli with a 10Sa RNA decapeptide, J Biol Chem 270, 9322–9326.

13. Gottesman, S., Roche, E., Zhou, Y., and Sauer, R. T. (1998) The ClpXP and ClpAP proteases degrade proteins with carboxy-terminal peptide tails added by the SsrA-tagging system, Genes Dev 12, 1338–1347.

14. Gottesman, S. (1996) Proteases and their targets in Escherichia coli, Annu Rev Genet 30, 465–506.

15. Sauer, R. T., and Baker, T. A. (2011) AAA+ proteases: ATP-fueled machines of protein destruction, Annu Rev Biochem 80, 587–612.

16. Olivares, A. O., Baker, T. A., and Sauer, R. T. (2018) Mechanical Protein Unfolding and Degradation, Annu Rev Physiol 80, 413–429.

17. Horwich, A. L., Weber-Ban, E. U., and Finley, D. (1999) Chaperone rings in protein folding and degradation, Proc Natl Acad Sci U S A 96, 11033–11040.

18. Olivares, A. O., Baker, T. A., and Sauer, R. T. (2016) Mechanistic insights into bacterial AAA+ proteases and protein-remodelling machines, Nat Rev Microbiol 14, 33–44.

19. Maillard, R. A., Chistol, G., Sen, M., Righini, M., Tan, J., Kaiser, C. M., Hodges, C., Martin, A., and Bustamante, C. (2011) ClpX(P) generates mechanical force to unfold and translocate its protein substrates, Cell 145, 459–469.

20. Bell, T. A., Baker, T. A., and Sauer, R. T. (2019) Interactions between a subset of substrate side chains and AAA+ motor pore loops determine grip during protein unfolding, eLife 8.

21. Lin, L., and Ghosh, S. (1996) A glycine-rich region in NF-kappaB p105 functions as a processing signal for the generation of the p50 subunit, Mol Cell Biol 16, 2248–2254.

22. Too, P. H., Erales, J., Simen, J. D., Marjanovic, A., and Coffino, P. (2013) Slippery substrates impair function of a bacterial protease ATPase by unbalancing translocation versus exit, J Biol Chem 288, 13243–13257.

23. Martin, A., Baker, T. A., and Sauer, R. T. (2008) Pore loops of the AAA+ ClpX machine grip substrates to drive translocation and unfolding, Nat Struct Mol Biol 15, 1147–1151.

24. Flynn, J. M., Levchenko, I., Seidel, M., Wickner, S. H., Sauer, R. T., and Baker, T. A. (2001) Overlapping recognition determinants within the ssrA degradation tag allow modulation of proteolysis, Proceedings of the National Academy of Sciences of the United States of America 98, 10584–10589.

25. Levchenko, I., Seidel, M., Sauer, R. T., and Baker, T. A. (2000) A specificity-enhancing factor for the ClpXP degradation machine, Science 289, 2354–2356.

26. Neher, S. B., Villén, J., Oakes, E. C., Bakalarski, C. E., Sauer, R. T., Gygi, S. P., and Baker, T. A. (2006) Proteomic profiling of ClpXP substrates after DNA damage reveals extensive instability within SOS regulon, Mol Cell 22, 193–204.

27. Pruteanu, M., and Baker, T. A. (2009) Controlled degradation by ClpXP protease tunes the levels of the excision repair protein UvrA to the extent of DNA damage, Mol Microbiol 71, 912–924.

28. Neher, S. B., Flynn, J. M., Sauer, R. T., and Baker, T. A. (2003) Latent ClpX-recognition signals ensure LexA destruction after DNA damage, Genes Dev 17, 1084–1089.

29. Camberg, J. L., Hoskins, J. R., and Wickner, S. (2009) ClpXP protease degrades the cytoskeletal protein, FtsZ, and modulates FtsZ polymer dynamics, Proc Natl Acad Sci U S A 106, 10614–10619.

30. Flynn, J. M., Neher, S. B., Kim, Y.-I., Sauer, R. T., and Baker, T. A. (2003) Proteomic Discovery of Cellular Substrates of the ClpXP Protease Reveals Five Classes of ClpX-Recognition Signals, Molecular Cell 11, 671–683.

31. McGinness, K. E., Baker, T. A., and Sauer, R. T. (2006) Engineering controllable protein degradation, Mol Cell 22, 701–707.

32. Davis, J. H., Baker, T. A., and Sauer, R. T. (2011) Small-Molecule Control of Protein Degradation Using Split Adaptors, Acs Chemical Biology 6, 1205–1213.

33. Davis, J. H. (2010) Understanding and Harnessing Energy-Dependent Proteolysis for Controlled Protein Degredation in Bacteria, Thesis - MIT.

34. Jiang, T., Xing, B., and Rao, J. (2008) Recent developments of biological reporter technology for detecting gene expression, Biotechnol Genet Eng Rev 25, 41–75.

35. Cormack, B. P., Valdivia, R. H., and Falkow, S. (1996) FACS-optimized mutants of the green fluorescent protein (GFP), Gene 173, 33–38.

36. Guzman, L. M., Belin, D., Carson, M. J., and Beckwith, J. (1995) Tight regulation, modulation, and high-level expression by vectors containing the arabinose PBAD promoter, J Bacteriol 177, 4121–4130.

37. Siegele, D. A., and Hu, J. C. (1997) Gene expression from plasmids containing the araBAD promoter at subsaturating inducer concentrations represents mixed populations, Proceedings of the National Academy of Sciences of the United States of America 94, 8168–8172.

38. Warren, J. W., Walker, J. R., Roth, J. R., and Altman, E. (2000) Construction and characterization of a highly regulable expression vector, pLAC11, and its multipurpose derivatives, pLAC22 and pLAC33, Plasmid 44, 138–151.

39. Lutz, R., and Bujard, H. (1997) Independent and tight regulation of transcriptional units in Escherichia coli via the LacR/O, the TetR/O and AraC/I1-I2 regulatory elements, Nucleic Acids Res 25, 1203–1210.

40. Li, G. W., Burkhardt, D., Gross, C., and Weissman, J. S. (2014) Quantifying absolute protein synthesis rates reveals principles underlying allocation of cellular resources, Cell 157, 624–635.

41. Lederer, T., Kintrup, M., Takahashi, M., Sum, P. E., Ellestad, G. A., and Hillen, W. (1996) Tetracycline analogs affecting binding to Tn10-Encoded Tet repressor trigger the same mechanism of induction, Biochemistry 35, 7439–7446.

42. Gur, E., and Sauer, R. T. (2008) Evolution of the ssrA degradation tag in Mycoplasma: specificity switch to a different protease, Proc Natl Acad Sci U S A 105, 16113–16118.

43. Cameron, D. E., and Collins, J. J. (2014) Tunable protein degradation in bacteria, Nature biotechnology 32, 1276–1281.

44. Reis, A. C., and Salis, H. M. (2020) An Automated Model Test System for Systematic Development and Improvement of Gene Expression Models, ACS Synthetic Biology 9, 3145–3156.

45. Miller, J. H. (1972) Experiments in molecular genetics, Cold Spring Harbor Laboratory, [Cold Spring Harbor, N.Y.].

46. Chalfie, M., Tu, Y., Euskirchen, G., Ward, W. W., and Prasher, D. C. (1994) Green fluorescent protein as a marker for gene expression, Science 263, 802–805.

47. Martin, A., Baker, T. A., and Sauer, R. T. (2008) Protein unfolding by a AAA+ protease is dependent on ATP-hydrolysis rates and substrate energy landscapes, Nat Struct Mol Biol 15, 139–145.

48. Kenniston, J. A., Baker, T. A., and Sauer, R. T. (2005) Partitioning between unfolding and release of native domains during ClpXP degradation determines substrate selectivity and partial processing, Proc Natl Acad Sci U S A 102, 1390–1395.

49. Lee, C., Schwartz, M. P., Prakash, S., Iwakura, M., and Matouschek, A. (2001) ATP-dependent proteases degrade their substrates by processively unraveling them from the degradation signal, Mol Cell 7, 627–637.

50. Chen, X., Zaro, J. L., and Shen, W. C. (2013) Fusion protein linkers: property, design and functionality, Adv Drug Deliv Rev 65, 1357–1369.

51. Andersen, J. B., Sternberg, C., Poulsen, L. K., Bjorn, S. P., Givskov, M., and Molin, S. (1998) New unstable variants of green fluorescent protein for studies of transient gene expression in bacteria, Appl Environ Microbiol 64, 2240–2246.

52. Kim, Y. I., Burton, R. E., Burton, B. M., Sauer, R. T., and Baker, T. A. (2000) Dynamics of substrate denaturation and translocation by the ClpXP degradation machine, Mol Cell 5, 639–648.

53. Bohn, C., Binet, E., and Bouloc, P. (2002) Screening for stabilization of proteins with a trans-translation signature in Escherichia coli selects for inactivation of the ClpXP protease, Mol Genet Genomics 266, 827–831.

54. Wah, D. A., Levchenko, I., Baker, T. A., and Sauer, R. T. (2002) Characterization of a specificity factor for an AAA+ ATPase: assembly of SspB dimers with ssrA-tagged proteins and the ClpX hexamer, Chem Biol 9, 1237–1245.

55. Farrell, C. M., Grossman, A. D., and Sauer, R. T. (2005) Cytoplasmic degradation of ssrA-tagged proteins, Mol Microbiol 57, 1750–1761.

56. Lewis, M. (2005) The lac repressor, C R Biol 328, 521–548.

57. Szydlo, K., Ignatova, Z., and Gorochowski, T. E. (2022) Improving the Robustness of Engineered Bacteria to Nutrient Stress Using Programmed Proteolysis, ACS Synth Biol 11, 1049–1059.

58. Douglass, E. F., Jr., Miller, C. J., Sparer, G., Shapiro, H., and Spiegel, D. A. (2013) A comprehensive mathematical model for three-body binding equilibria, J Am Chem Soc 135, 6092–6099.

59. Erlich, H. A., Cohen, S. N., and McDevitt, H. O. (1978) A sensitive radioimmunoassay for detecting products translated from cloned DNA fragments, Cell 13, 681–689.

60. Wah, D. A., Levchenko, I., Rieckhof, G. E., Bolon, D. N., Baker, T. A., and Sauer, R. T. (2003) Flexible linkers leash the substrate binding domain of SspB to a peptide module that stabilizes delivery complexes with the AAA+ ClpXP protease, Mol Cell 12, 355–363.

61. 61. Hersch, G. L., Baker, T. A., and Sauer, R. T. (2004) SspB delivery of substrates for ClpXP proteolysis probed by the design of improved degradation tags, Proc Natl Acad Sci U S A 101, 12136–12141.

62. Jacobson, R. H., Zhang, X. J., DuBose, R. F., and Matthews, B. W. (1994) Three-dimensional structure of beta-galactosidase from E. coli, Nature 369, 761–766.

63. Jacob, F., and Monod, J. (1961) Genetic regulatory mechanisms in the synthesis of proteins, J Mol Biol 3, 318–356.

64. Newton, W. A., Beckwith, J. R., Zipser, D., and Brenner, S. (1965) Nonsense mutants and polarity in the lac operon of Escherichia coli, J Mol Biol 14, 290–296.

65. Goldschmidt, R. (1970) In vivo degradation of nonsense fragments in E. coli, Nature 228, 1151–1154.

66. Rickenberg, H. V., Yanofsky, C., and Bonner, D. M. (1953) Enzymatic deadaptation, J Bacteriol 66, 683–687.

67. Kuzminov, A. (1999) Recombinational repair of DNA damage in Escherichia coli and bacteriophage lambda, Microbiol Mol Biol Rev 63, 751–813, table of contents.

68. Datsenko, K. A., and Wanner, B. L. (2000) One-step inactivation of chromosomal genes in Escherichia coli K-12 using PCR products, Proceedings of the National Academy of Sciences of the United States of America 97, 6640–6645.

69. Mahmood, T., and Yang, P. C. (2012) Western blot: technique, theory, and trouble shooting, N Am J Med Sci 4, 429–434.

70. Towbin, H., Staehelin, T., and Gordon, J. (1979) Electrophoretic transfer of proteins from polyacrylamide gels to nitrocellulose sheets: procedure and some applications, Proc Natl Acad Sci U S A 76, 4350–4354.

71. Jacobson, M. K., and Lark, K. G. (1973) DNA replication in Escherichia coli: evidence for two classes of small deoxyribonucleotide chains, J Mol Biol 73, 371–396.

72. Bochner, B. R., and Ames, B. N. (1982) Complete analysis of cellular nucleotides by two-dimensional thin layer chromatography, J Biol Chem 257, 9759–9769.

73. Rogers, A. T., Bullard, K. R., Dod, A. C., and Wang, Y. (2022) Bacterial Growth Curve Measurements with a Multimode Microplate Reader, Bio Protoc 12, e4410.

