## Supporting Information for "Degron-controlled protein degradation in *Escherichia coli*: New Approaches and Parameters"

### Table of Contents

**Supplemental Figures**

Figure S1. C-terminal fusion of SsrA or LAA tags to SspB........ S3

Figure S2. Complementation of Δ*clpX* restores normal SspB-sensitivity of recA-DAS#2 and ruvB-DAS#2 degron alleles........ S4

Figure S3. Effect of long linker regions on SspB-dependent degradation of GFP-DAS........ S5

Figure S4. UV-resistance of *recA*- and *ruvB*-DAS#3........ S6

Figure S5. Near constitutive expression of *GFP* integrated at the *lacZ* locus........ S7

Figure S6. Background-subtracted GFP fluorescence of lacZ::GFP-DAS#3 and pGFP-DAS#3........ S8

Figure S7. Comparison of relative GFP expression from high copy plasmid under control of the lacIq promoter, and a genome resident *GFP* cloned in place of the *lacZ* coding sequence........ S9

Figure S8. Growth curves for all strains and growth conditions from Figure 7........ S10

Figure S9. Titration of the inducer anhydrotetracycline (aTc)........ S11

Figure S10. Genome integration of the *sspB-rs* expression cassette at the *sspB* loci results in significant leakiness of SspB-rs expression........ S12

**Supplementary Tables**

Table S1. Strains used in this work........ S13

Table S2. Plasmids used in this work........ S15

Table S3. Primers and design details........ S16

**References**........ S25

#### Supplementary Figures

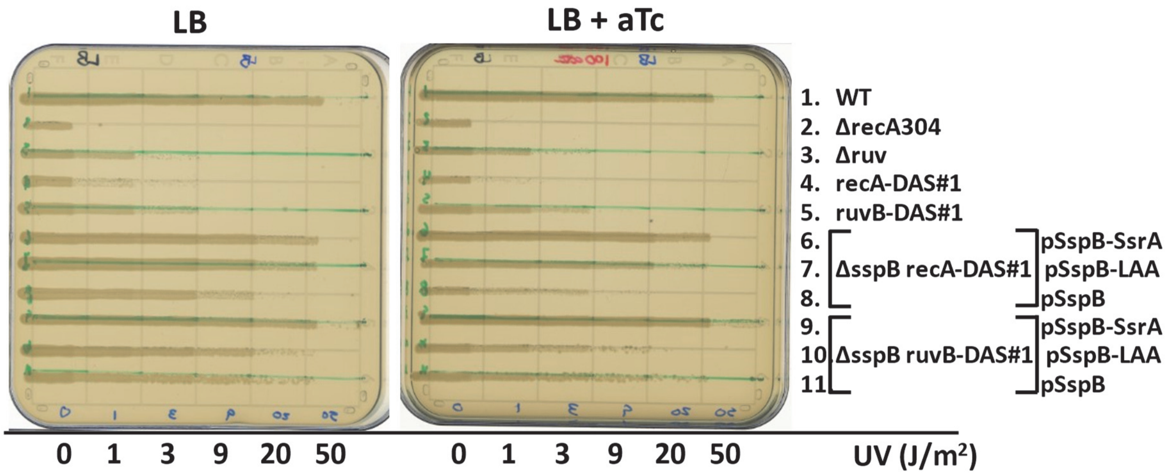

**Figure S1. C-terminal fusion of SsrA or LAA tags to SspB.**

In this figure, a similar plasmid backbone, differing only in the ribosome binding site from that used for pSspB and pSspB-rs was used for SspB expression. Plasmids contained either unmodified *sspB* (pSspB), or C-terminal *sspB* fusions with either wild-type SsrA sequence (AANDENYALAA, pSspB-SsrA) or C-terminal SsrA ClpX-interacting motif (LAA, pSspB-LAA). From single colonies on selective plates, cultures were grown overnight at 28 °C in LB+Cm (plasmid-carrying strains) or LB (no plasmid) and the next day diluted 1:100 into the same media and grown at 28 °C to mid-log phase ($OD_{600}$ = 0.2 – 0.5). Portions of each culture were then deposited in a horizontal line by drawing across the plate using a capillary tube. Once the plates had dried, a masking method was employed to deliver the indicated cumulative UV doses in vertical stripes, and the plates were incubated at 28 °C overnight in the dark before imaging. Note that the plates do not contain selective antibiotic, which may explain the appearance of colony-size variation observed in lanes 10 and 11. Where indicated, anhydrotetracycline (aTc) was added to a final 1x concentration of 100 µg/mL. WT is AB1157, Δ*recA304* is JC10287, Δ*ruv* is AM547, *recA*-DAS#1 is eGC058, Δ*sspB recA*-DAS#1 is eGC059, *ruvB*-DAS#1 is eGC036, Δ*sspB ruvB*-DAS#1 is eGC061, pSspB is pGC336, pSspB-SsrA is pGC332, pSspB-LAA pGC334.

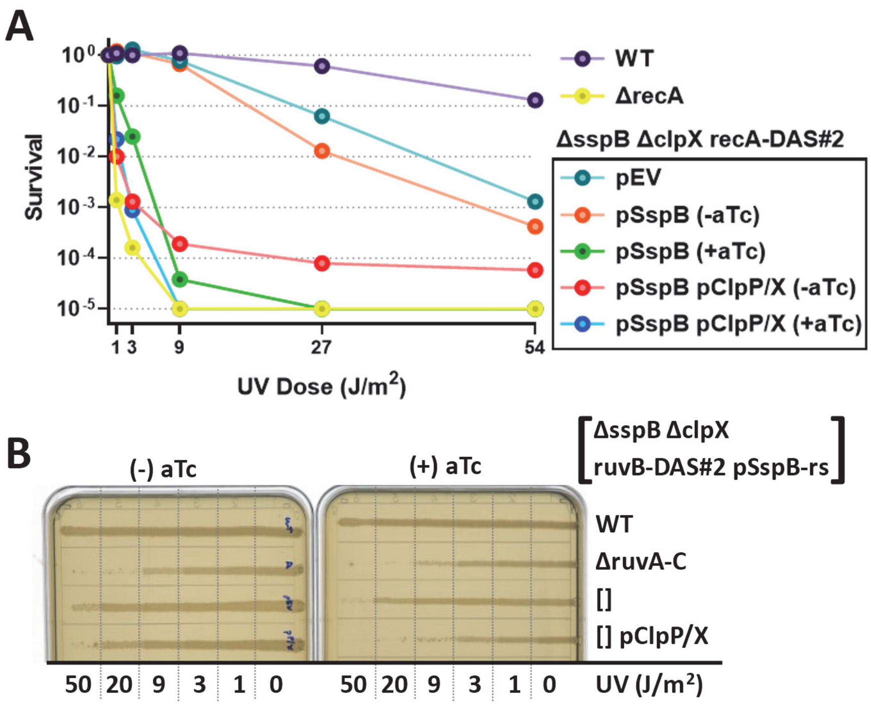

**Figure S2**. **Complementation of Δ*clpX* restores normal SspB-sensitivity of recA-DAS#2 and ruvB-DAS#2 degron alleles.**

**A.** The background for all strains is AB1157. All strains contain two plasmids, either the indicated SspB- or ClpP/X-containing plasmid(s) or empty vector(s). pSspB is the leaky SspB plasmid, lacking the heterologous *rs* (reduced stability) element. pClpP/X contains the *clpP-clpX* operon under control of the native promoter, where 157 bp of sequence 5' to *clpP* was cloned in place of the hybrid lac-ara promoter of pZS*24-MCS (1). From a saturated overnight culture, 1:100 dilutions were made into LB + Cm, Km, and cultures grown at 28 °C to $OD_{600}$ = 0.2 - 0.5 (approximately 3 hours). Each strain was then spotted in 10-fold serial dilutions on LB agar plates containing the same antibiotics and with or without aTc, as indicated. Individual plates were exposed to the indicated dose of ultraviolet radiation (UV), incubated overnight at 28 °C in the dark, and survival was calculated as the observed titre normalized to unirradiated control. **B.** MG1655 pZA31-luc (WT), AM547 pZA31-luc (AB1157 Δ*ruv*), AB1157 ΔsspB recA-DAS#2 (eGC660) pSspB-rs (pGC533-1), and eGC660 pSspB-rs pClpP/X (pGC799-1) were scraped from fresh colonies and grown to $OD_{600}$=0.2 – 0.6 in LB + Cm (+ Km for pClpP/X strain) at 28 °C (~ three hours), and portions of each culture spread across two plates by capillary tube. Sections of each plate (delineated by the vertical dotted lines) were exposed to the cumulative doses of UV indicated and imaged after growth overnight at 28 °C in the dark.

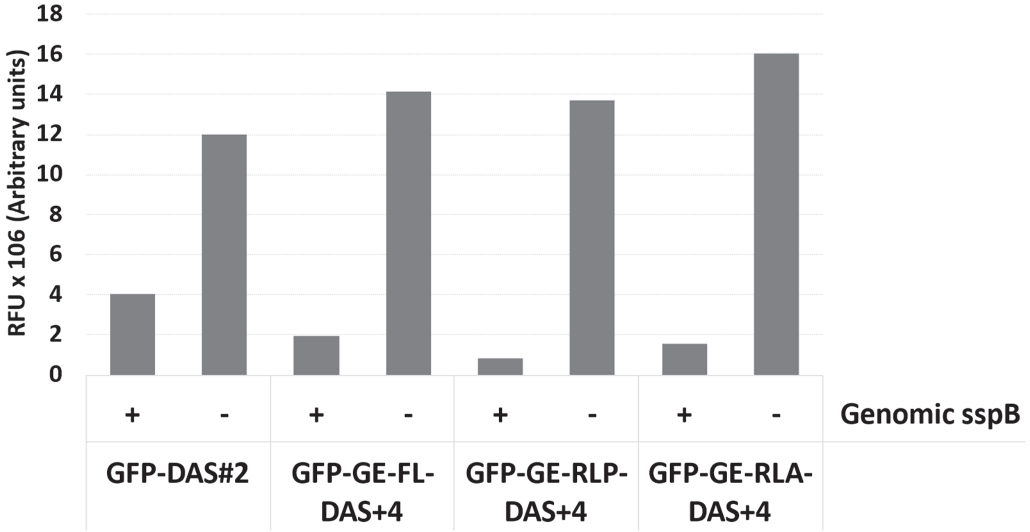

**Figure S3. Effect of long linker regions on SspB-dependent degradation of GFP-DAS.**

GE is the "Acidic" grip sequence (GEGDGEGDGEGD) from Bell et al. (2). FL is Flexible Linker (GGGSGGGGSGGGS), RLP is Rigid Linker Proline (PAPAPAPAPAPAPAPAPAPA), and RLA is Rigid Linker Alanine (AEAAAKEAAAKEAAAKEAAAKALEAEAAAKEAAAKEAAAKA) summarized in Chen et al. (3). DAS+4 is the SspB-specific degron sequence (AANDENYSENYADAS) from McGinness et al. (4). The GFP employed is the mut3b variant, C-terminally truncated as in Bell et al (2). Data were normalized to subtract the background fluorescence of the empty vector containing WT strain. For this experiment, from a saturated overnight culture cells were diluted 1:285 in 2 mL of LB + Km and grown at 28 °C for 4 hours to a final $OD_{600}$ = 0.4 – 0.5. The entire two mL of culture was pelleted, washed 1x in PBS, fixed in 1mL of 4% paraformaldehyde in PBS + 2 mM PIPES pH 7.2 for 15 min at room temperatures, and then washed 2x with 1 mL PBS. Then, $4 \times 10^8$ fixed cells were pelleted, resuspended in 20 uL of PBS and spotted directly on the fluor stage of a Typhoon FLA 7000 laser scanner (GE Biosciences) and the GFP fluorescence measured by excitation with the 473 nm laser and emission signal collected through the 520 long-pass filter. MG1655 (-) or eGC027 (+) carrying one of: pGFP-GE-FL-DAS+4 (pGC639), pGFP-GE-RLP-DAS+4 (pGC640) or pGFP-GE-RLA-DAS+4 (pGC643).

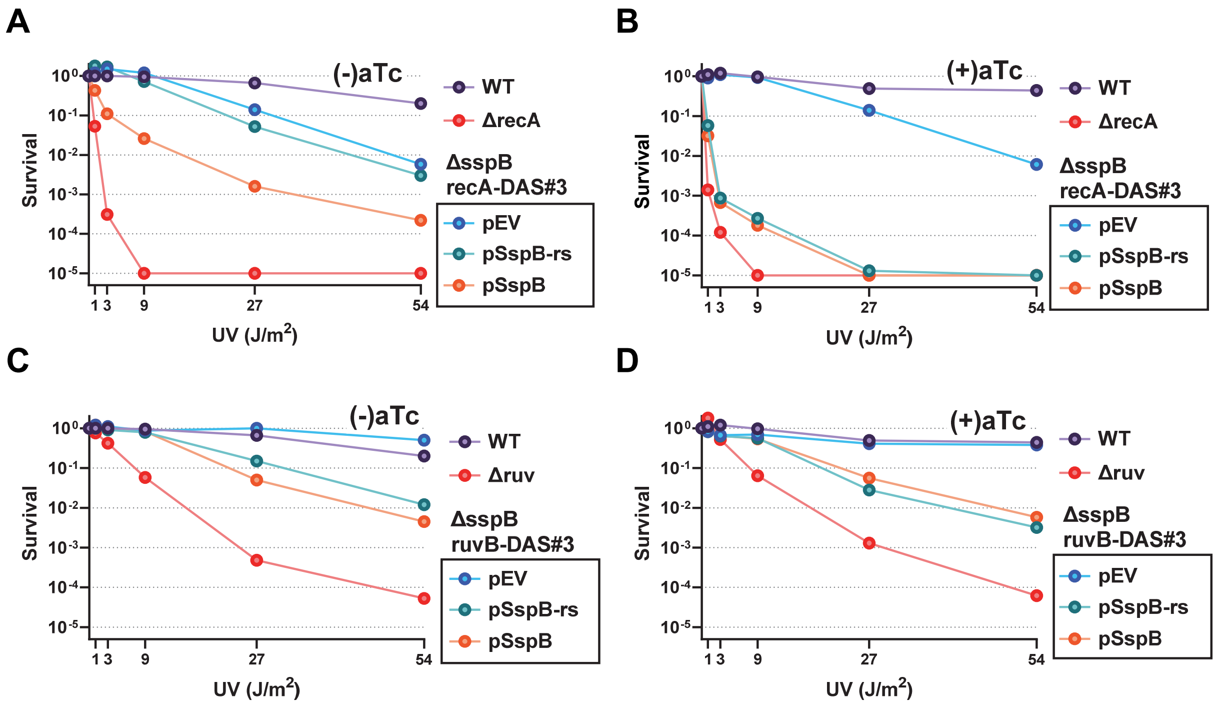

**Figure S4. UV-resistance of recA- and ruvB-DAS#3.**

From a saturated overnight culture, 1:100 dilutions were made into LB + Cm, and cultures grown at 28 °C to $OD_{600}$ = 0.2 - 0.5 (approximately 3 hours). Each strain was then spotted in 10-fold serial dilutions on LB agar plates containing Cm, with or without aTc, as indicated. Individual plates were exposed to the indicated dose of ultraviolet radiation (UV), incubated overnight at 28 °C in the dark, and survival calculated as the observed titer normalized to unirradiated control. WT is AB1157 pZA31-luc, Δ*ruv* is AM547 pZE31-luc, Δ*sspB recA*-DAS#3 is eGC698, Δ*sspB ruvB*-DAS#3 is eGC699, pEV is pZA31-luc, pSspB is pGC513-1 and pSspB-rs is pGC533-1.

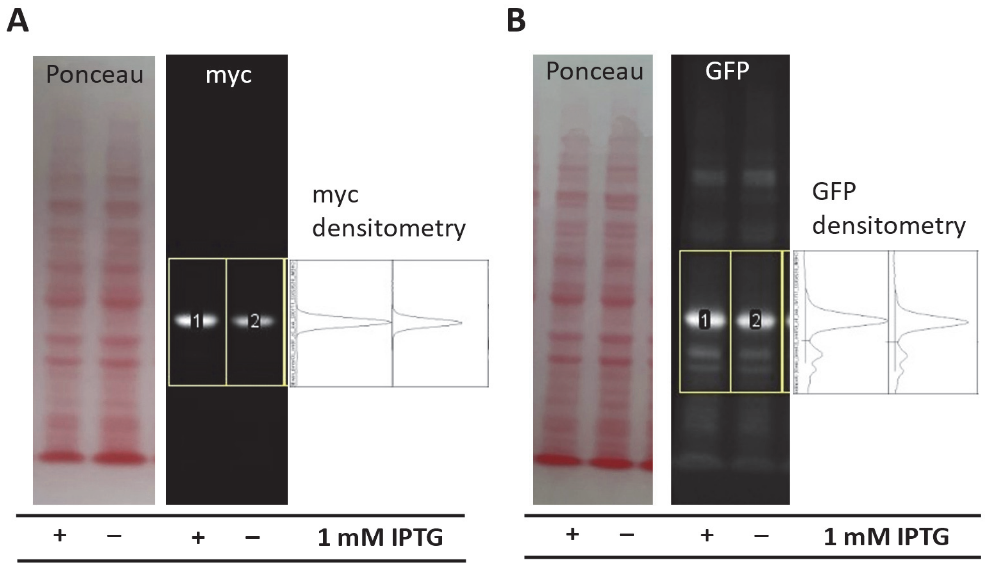

C

MG1655 ΔsspB ΔlacZ::GFP-DAS#3
(+) IPTG vs (-) IPTG fold enrichment

| | |
|---|---|
| myc signal | 1.4 x |
| GFP signal | 1.2 x |

**Figure S5. Near constitutive expression of *GFP* integrated at the *lacZ* locus.**

The precise replacement of the *lacZ* coding sequence with that of GFP-DAS#3 in the MG1655 Δ*sspB* background. This experiment was structured as given for the "Beta-galactosidase western blot" section in Materials and Methods, with these samples being harvested as T = 0 timepoints (before tetracycline addition) as in Figure 6 of the main text. **A.** Ponceau S and α-myc western blot signals of the same membrane, with densitometry of the western signal for the MG1655 Δ*sspB* Δ*lacZ*::GFP-DAS#3 strain grown in the presence or absence of 1 mM IPTG. **B.** A replicate blot for the same samples as in panel A, but probed with α-GFP. **C.** The apparent fold increase of myc and GFP western blot signals due to the presence of 1 mM IPTG.

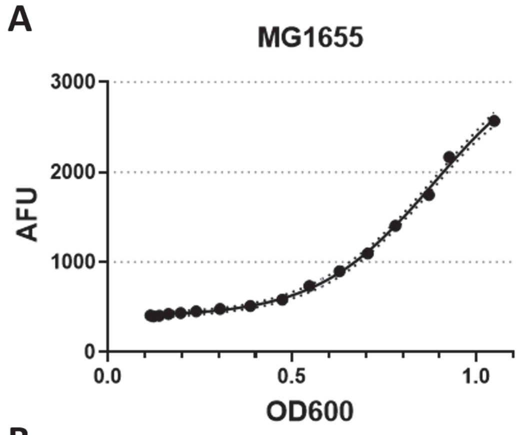

B

| Adjusted R squared | 0.9972 |
|---|---|
| Best-fit values | |
| Top | 3362 |
| Bottom | 393.0 |
| LogIC50 | 0.8834 |
| HillSlope | 2.762 |
| IC50 | 7.645 |
| Span | 2969 |

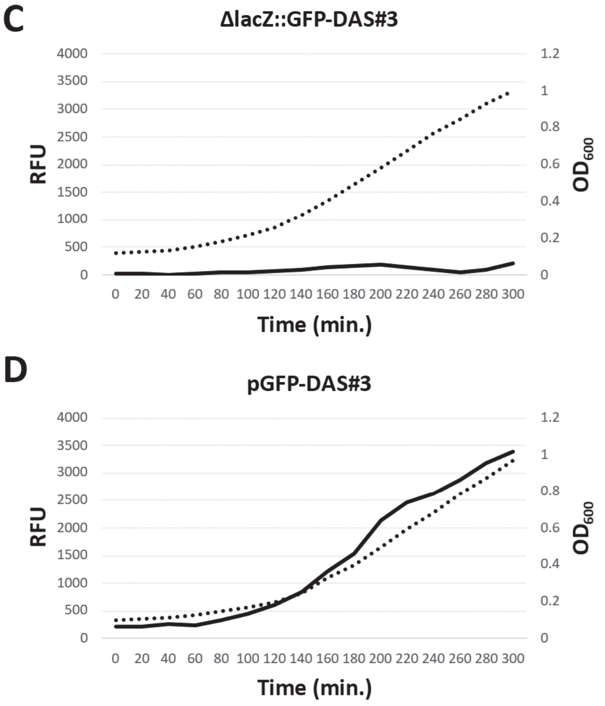

**Figure S6. Background-subtracted GFP fluorescence of lacZ::GFP-DAS#3 and pGFP-DAS#3.**

**A.** Background GFP (auto) fluorescence versus $OD_{600}$ for the MG1655 WT strain (large dots), and best fit for a four parameter sigmoid with variable slope (solid line) including 95% confidence interval (dotted lines). **B.** Parameters and goodness of fit for the fitted curve from Panel A. Panels **C and D**, OD600 (dotted line) and background-subtracted GFP signal (solid line) for the MG1655 Δ*sspB* Δ*lacZ*::GFP-DAS#3 (Panel C) and MG1655 Δ*sspB* pGFP-DAS#3 (Panel D) strains. Data were collected during growth in MOPS, 0.2% CAA, 0.2 % glucose, plus Km, at 37 °C after a 1:50 dilution from overnight cultures grown in LB. Data were collected in a plate reader, as described in Materials and Methods, except with gain = 40. The WT strain is eGC706, Δ*lacZ*::GFP-DAS#3 is eGC762, pGFP-DAS#3 is MG1655 pGC707-1.

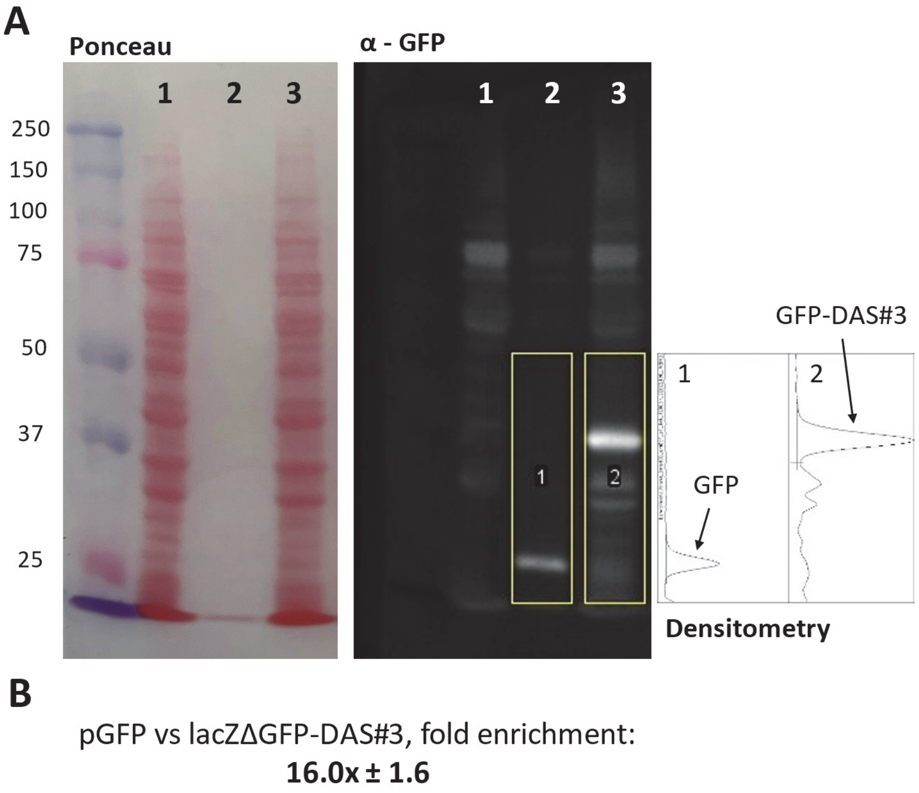

**Figure S7. Comparison of relative GFP expression from high copy plasmid under control of the lacIq promoter, and a genome resident GFP cloned in place of the lacZ coding sequence.**

**A.** Representative α-GFP western blot showing lane 1 - WT (no GFP), lane 2 – MG1655 pZE27GFP (“pGFP”), lane 3 – MG1655 Δ*sspB* lacZΔ::GFP-DAS#3 (eGC706, “lacZΔGFP-DAS#3”). Ponceau S stained blot at left, GFP signal at middle. Note that the $OD_{600}$ adjusted pGFP lysate was diluted 1:50 before loading. At left, densitometric 1-D lane plots, bounded by the yellow boxes in the middle panel, showing the enclosed curve areas used for quantification in ImageJ. **B.** The mean and SEM fold excess of GFP signal (pGFP compared to lacZΔ::GFP) of three independent measurements, n = 3. All growth in LB, including Km for pGFP and 1 mM IPTG for lacZΔ::GFP-DAS#3. For the images above, from a saturated overnight culture a 1:100 dilution was made, and the subcultures incubated at 37 °C for 2.5 hours to OD600 of 0.8 - 0.9, cultures were again back-diluted into fresh LB this time to a calculated $OD_{600}$ = 0.00625 and grown at 37 °C for seven doublings to $OD_{600}$ = 0.75 – 0.95. Pellets equal to 1.5 ODs of cells were used to produce crude lysates for western blot (Materials and Methods). For the n = 2 repetition a similar experiment was performed but after the second back-dilution cultures were grown for two doublings to a similar final OD600. For n =3, no second back-dilution was performed, and cells were harvested at OD600 = 0.6 – 0.8.

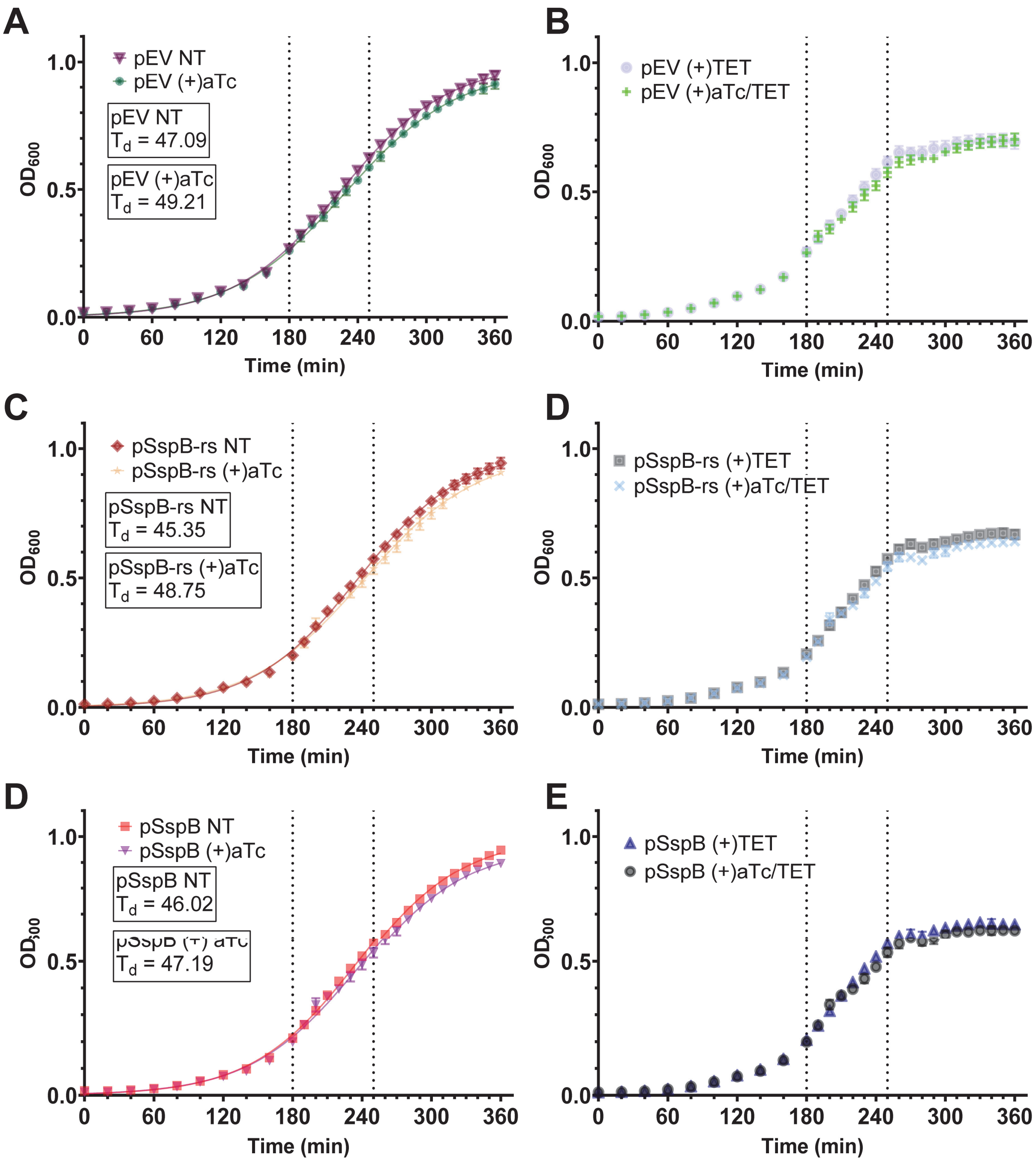

**Figure S8. Growth curves for all strains and growth conditions from Figure 7.**

Growth curve data (optical density at 600 nm, $OD_{600}$) collected during the GFP fluorescence data given in the experiments of Figure 7. See the main text for experimental details. The vertical dotted lines indicate the time of anhydrotetracycline addition (aTc, added to a final concentration of 100 ng/mL at 180 minutes) or tetracycline (TET, added to a final concentration of 10 µg/mL at 250 minutes). Note that aTc and TET were not added in all cases; their addition is denoted in the sample name in each panel. Doubling times ($T_d$, in minutes) were calculated from the logistic growth equation in Graphpad Prism.

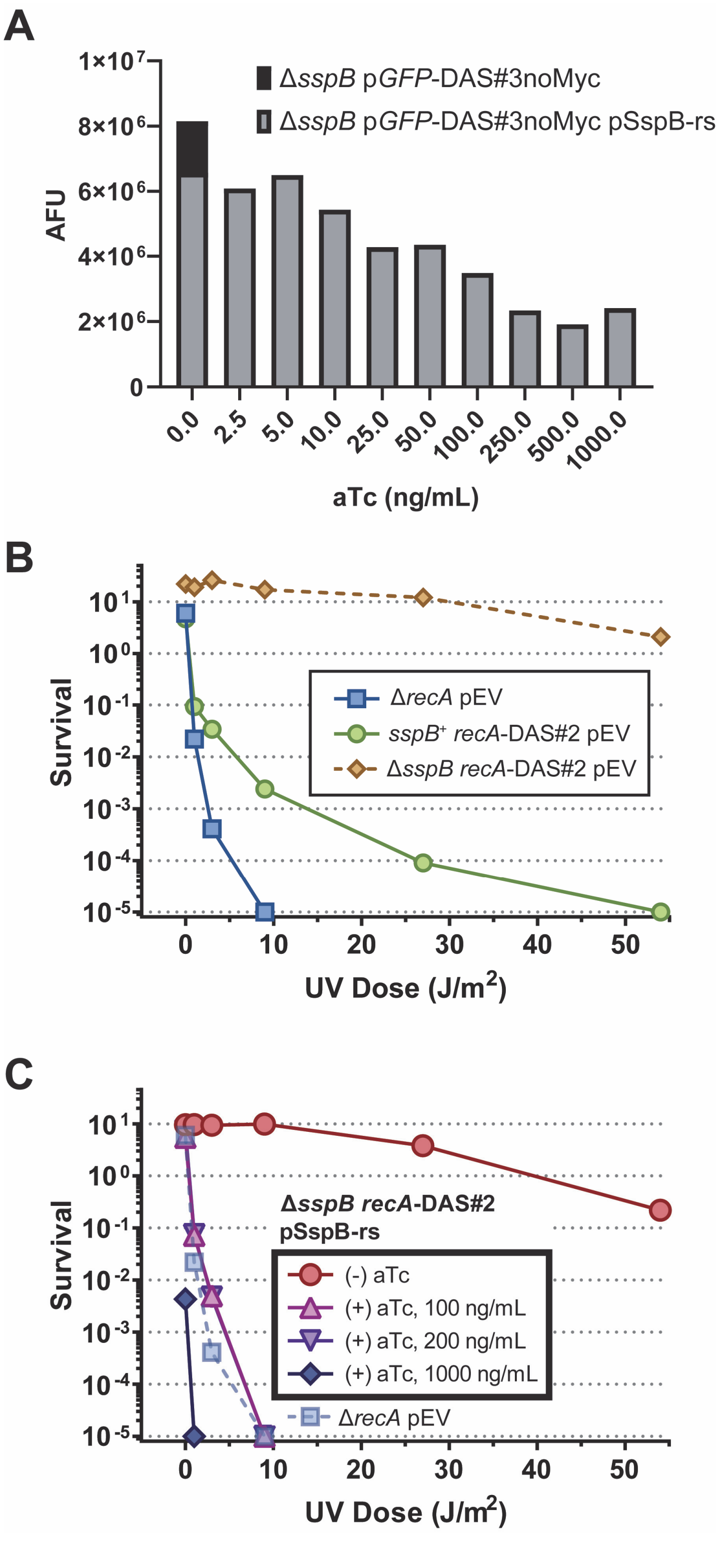

**Figure S9. Titration of the inducer anhydrotetracycline (aTc).**

**A.** GFP fluorescence (Arbitrary fluorescence units, AFU) measured during exponential growth of strains containing the degron-tagged GFP from Figure 7 (GFP-DAS#3noMyc construct, see Primers for the DAS#3 sequence). Cells were grown overnight to saturation in LB media containing kanamycin (Δ*sspB* p*GFP*-DAS#3noMyc) or kanamycin and chloramphenicol (Δ*sspB* p*GFP*-DAS#3noMyc pSspB-rs) at 28°C. The overnight cultures diluted 1:300 into fresh LB containing the same antibiotics, and incubated at 28°C with vigorous aeration to $OD_{600}$ = 0.3 - 0.4. One milliliter of culture was pelleted and subjected to the fixation protocol given in Figure S3. The density of the fixed cells was determined by $OD_{600}$ and 2 x $10^8$ cells (Equivalent to 1 mL of $OD_{600}$ = 0.1) were pelleted, resuspended in 20 uL and spotted on the fluor stage of a Typhoon FLA 7000 for quantification of GFP (see Figure S3). All measurements are background subtracted for MG1655 wild-type, without GFP. MG1655 is the background for both strains: Δ*sspB* is eGC027, p*GFP*-DAS#3noMyc is pGC643-1 and pSspB-rs is pGC533-1. **B and C.** UV survival assay, performed in a single experiment but split into two panels for clarity. The experiment used the standard assay conditions provided in Materials and Methods. However, unlike the other UV-survival figures, here the data were not normalized to the unirradiated control to make clear the loss of titre when plating on LB agar plates containing 1000 ng/mL of aTc. The Δ*recA* survival curve is duplicated between panels. Strains are in the AB1157 background: Δ*recA* is JC10287, sspB$^+$ recA-DAS#2 is eGC566, ΔsspB recA-DAS#2 is eGC567, pEV is pZA31-luc, and pSspB-rs is pGC533-1.

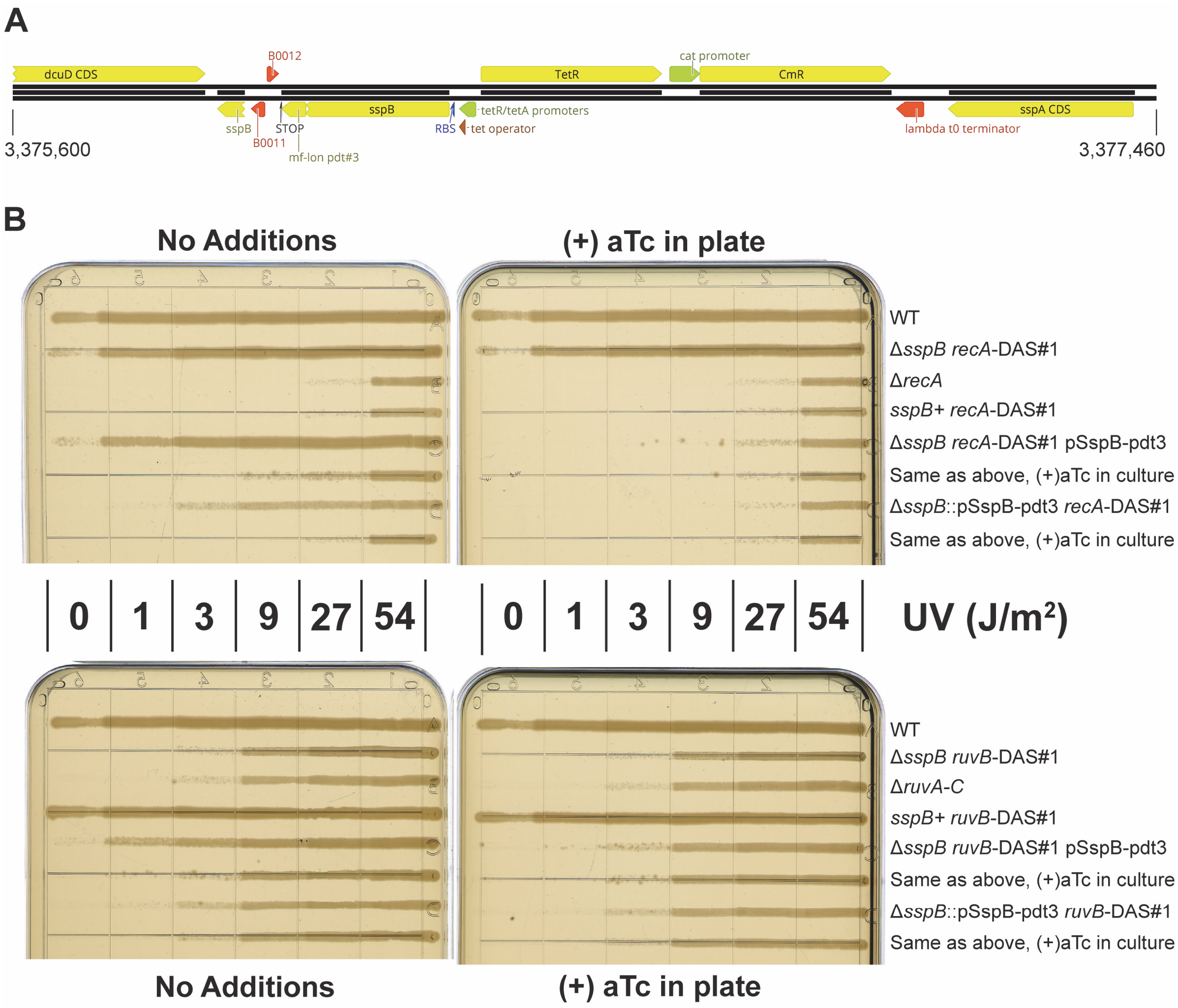

**Figure S10. Genome integration of the *sspB*-rs expression cassette at the *sspB* loci results in significant leakiness of SspB-rs expression.**

**A.** Structure of the genome integrations tested in panel B. All plasmid sequence, except for that required for replication, was integrated at the *sspB* loci by homology-directed integration (5), as depicted. The coordinates provided are for MG1655 (NCBI Reference Sequence: NC_000913.3). **B.** Qualitative UV-survival assay. Strains were grown overnight in LB at 28°C, plus chloramphenicol for strains containing pSspB-pdt3, to saturation. Overnight cultures were diluted 1:100 into the same media with duplicate cultures for pSspB-pdt3 either with, or without, aTc. Cultures were incubated at 28°C until a cell density of $OD_{600}$ = 0.2 – 0.4, at which point portions of the culture were spread across plates with a capillary tube. Immediately after streaks dried plates were irradiated at the indicated UV doses. Plates were photographed following 18 hours of incubation in the dark at 28°C. Strains are in the AB1157 background; WT is AB1157, Δ*recA* is JC10287, Δ*ruvA-C* is AM547, *sspB*$^{+}$ *recA*-DAS#1 is eGC058, Δ*sspB* *recA*-DAS#1 is eGC059, Δ*sspB*::pSspB-pdt3 *recA*-DAS#1 is eGC570, *sspB*$^{+}$ *ruvB*-DAS#1 is eGC036, Δ*sspB* *ruvB*-DAS#1 is eGC038, Δ*sspB*::pSspB-pdt3 *ruvB*-DAS#1 is eGC571, pSspB-pdt3 is pGC514-1. The pdt3 sequence is "PDT#3" – AANKNEENTNEVPTFMLNAGQANRRRV, from Cameron and Collins, 2014 (6).

### 1. Supplementary Tables

**Table S1.** Strains used in this work.

| Strain | Background | Genotype or description | Reference |
|---|---|---|---|
| **Published** | | | |
| MG1655[a] | MG1655 | Wild Type | Bachmann, B. J. (1972). Bacteriological Reviews 36(4): 525-557. |
| AB1157[b] | AB1157 | Wild Type | Bachmann, B. J. (1972). Bacteriological Reviews 36(4): 525-557. |
| JW3197-1 | Δ*sspB::kan* | CSGC #10426 | Baba, T., et al. (2006). Mol Syst Biol 2: 2006 0008. |
| JC10287 | AB1157 | Δ*(srlR-recA)304* | Csonka, L. N. and A. J. Clark (1979). Genetics 93(2): 321-343. |
| AM547 | AB1157 | Δ*(ruvA-ruvC)65* | Mandal, T. N., et al. (1993). J Bacteriol 175(14): 4325-4334. |
| JT33 | MG1655 | Δ*lacZY::cat* | Chakravartty, V. and J. E. Cronan (2012). J Bacteriol 194(5): 1113-1126. |

[a] Complete genotype of the background is F- *λ- ilvG rfb-50 rph-1*.
[b] Complete genotype of the background is F- *λ- rac thi-1 hisG4* Δ*(gpt-proA)62 argE3 thr-1 leuB6 kdgK51 rfbD1 araC14 lacY1 galK2 xylA5 mtl-1 tsx-33 supE44(glnV44) rpsL31 (strR)*.

| Strain | Background | Genotype or description | Reference |
|---|---|---|---|
| **This work** | | | |
| eGC025 | MG1655 | Δ*sspB::kan* | MG1655 x P1 JW3197-1 |
| eGC026 | AB1157 | Δ*sspB::kan* | AB1157 x P1 JW3197-1 |
| eGC027 | MG1655 | Δ*sspB* | eGC025 x pCP20 |
| eGC028 | AB1157 | Δ*sspB* | eGC026 x pCP20 |
| eGC023 | MG1655 | *recA-DAS#1-kan* | MG1655 x PCR pGC016 x pKD46 |
| eGC024 | MG1655 | *ruvB-DAS#1-kan* | MG1655 x PCR pGC016 x pKD46 |
| eGC054 | AB1157 | *recA-DAS#1-kan* | AB1157 x P1 eGC023 |
| eGC058 | AB1157 | *recA-DAS#1* | eGC054 x pCP20 |
| eGC055 | AB1157 | Δ*sspB recA-DAS#1-kan* | eGC028 x P1 eGC023 |
| eGC059 | AB1157 | Δ*sspB recA-DAS#1* | eGC055 x pCP20 |
| eGC036 | AB1157 | *ruvB-DAS#1-kan* | AB1157 x P1 eGC024 |
| eGC038 | AB1157 | Δ*sspB ruvB-DAS#1-kan* | eGC028 x P1 eGC024 |
| eGC061 | AB1157 | Δ*sspB ruvB-DAS#1* | eGC038 x pCP20 |
| eGC566 | AB1157 | *recA-DAS#2-kan* | AB1157 x PCR pGC561 x pKD46 |
| eGC567 | AB1157 | Δ*sspB recA-DAS#2-kan* | eGC028 x P1 eGC566 |
| eGC661 | AB1157 | Δ*sspB recA-DAS#2* | eGC567 x pCP20 |
| eGC568 | AB1157 | *ruvB-DAS#2-kan* | AB1157 x PCR pGC561 x pKD46 |
| eGC569 | AB1157 | Δ*sspB ruvB-DAS#2-kan* | eGC028 x P1 eGC568 |

| | | | |
|---|---|---|---|
| eGC570 | AB1157 | Δ*sspB*::pSspB-pdt3 *recA*-DAS#1 | eGC054 x PCR pGC533-1 x pKD46 |
| eGC571 | AB1157 | Δ*sspB*::pSspB-pdt3 *ruvB*-DAS#1 | eGC036 x PCR pGC533-1 x pKD46 |
| eGC663 | AB1157 | Δ*sspB ruvB-DAS#2* | eGC569 x pCP20 |
| eGC658 | MG1655 | Δ*clpX::kan* | MG1655 x PCR pKD13 x pDK46 |
| eGC668 | AB1157 | *recA-DAS#2 ΔclpX::kan* | eGC660 x P1 eGC658-1 |
| eGC669 | AB1157 | Δ*sspB recA-DAS#2 ΔclpX::kan* | eGC661 x P1 eGC658-1 |
| eGC670 | AB1157 | *ruvB-DAS#2 ΔclpX::kan* | eGC662 x P1 eGC658-1 |
| eGC671 | AB1157 | Δ*sspB ruvB-DAS#2* ΔclpX::kan | eGC663 x P1 eGC658-1 |
| eGC706 | MG1655 | Δ*sspB lacZ-DAS#3* | eGC027 x PCR pGC702 x pKD46 |
| eGC698 | MG1655 | Δ*sspB recA-DAS#3(no myc)-kan* | eGC027 x PCR pGC702 x pKD46 |
| eGC699 | MG1655 | Δ*sspB ruvB-DAS#3(no myc)-kan* | eGC027 x PCR pGC702 x pKD46 |
| eGC700 | MG1655 | Δ*sspB dnaG-DAS#2-kan* | eGC027 x PCR pGC561 x pKD46 |
| eGC697 | MG1655 | Δ*sspB holD-DAS#3(no myc)-kan* | eGC027 x PCR pGC561 x pKD46 |
| eGC607 | MG1655 | Δ*sspB ligA-DAS#2-kan* | eGC027 x PCR pGC561 x pKD46 |
| eGC693 | MG1655 | Δ*sspB ligA-DAS#3(no myc)-kan* | eGC027 x PCR pGC702 x pKD46 |
| eGC606 | MG1655 | Δ*sspB polA-DAS#2-kan* | eGC027 x PCR pGC561 x pKD46 |
| eGC665 | MG1655 | Δ*sspB polA-DAS#3(no myc)-kan* | eGC027 x PCR pGC702 x pKD46 |
| eGC610 | MG1655 | Δ*sspB priA-DAS#2-kan* | eGC027 x PCR pGC561 x pKD46 |
| eGC608 | MG1655 | Δ*sspB recB-DAS#2-kan* | eGC027 x PCR pGC561 x pKD46 |
| eGC666 | MG1655 | Δ*sspB recB-DAS#3(no myc)-kan* | eGC027 x PCR pGC702 x pKD46 |
| eGC609 | MG1655 | Δ*sspB recC-DAS#2-kan* | eGC027 x PCR pGC561 x pKD46 |
| eGC673 | MG1655 | *recC-DAS#3(no myc)-kan* | eGC004 x PCR pGC702 x pKD46 |
| eGC679 | MG1655 | Δ*sspB recC-DAS#3(no myc)-kan* | eGC027 x P1 eGC673 |
| L520 | MG1655 | Δ*sspB rnhA-DAS#3(no myc)-kan* | |

**Table S2.** Plasmids used in this work.

| Plasmid | Name | Replicon / Drug resistance / Other genes | Reference / Derivation |
|---|---|---|---|
| **Published** | | | |
| pKD4 | | R6K / *bla kan* | Datsenko, K. A. and B. L. Wanner (2000).<br>PNAS 97(12): 6640-6645. |
| pKD13 | | R6K / *bla kan* | Datsenko, K. A. and B. L. Wanner (2000).<br>PNAS 97(12): 6640-6645. |
| pKD46 | | SC101 / *bla* / *exo bet gam* | Datsenko, K. A. and B. L. Wanner (2000).<br>PNAS 97(12): 6640-6645. |
| pCP20 | | SC101 (ts) / *bla cat* / FLP | Cherepanov, P. P. and W. Wackernagel (1995).<br>Gene 158(1): 9-14. |
| pLAC22 | | ColE1 / *bla tet* / *lacI*$^{q}$ | Warren, J. W., et al. (2000).<br>Plasmid 44(2): 138-151. |
| pZA31-luc | | p15A / *cat* | Lutz, R. and H. Bujard (1997).<br>Nucleic Acids Res 25(6): 1203-1210. |
| pZS*24MCS | | pSC101* / *kan* | Lutz, R. and H. Bujard (1997).<br>Nucleic Acids Res 25(6): 1203-1210. |
| pdCas9-bacteria | | p15A / *cat* / tetR *cas9*(dead) | Addgene #44249.<br>Qi, L. S., et al. (2013).<br>Cell 152(5): 1173-1183. |
| pZE27GFP | | pBR322 / *kan* / GFPmut3b | Addgene #75452.<br>Cameron, D. E. and J. J. Collins (2014).<br>Nat Biotechnol 32(12): 1276-1281. |
| pCAS9-CR4 | | p15A / *cat* / tetR *cas9* | Addgene #62655.<br>Reisch, C. R. and K. L. Prather (2015).<br>Sci Rep 5: 15096. |
| **This work** | | | |
| pGC015 | | R6K / *bla kan* / (+) NheI | pKD4 |
| pGC016 | | R6K / *bla kan* / DAS#1 | pGC015 |
| pGC064 | pLAC22::sspB | ColE1 / *bla tet* / *lacI*$^{q}$ *sspB* | pLAC22 |
| pGC561-1 | | R6K / *bla kan* / DAS#2 | pGC015 |
| pGC513-1 | pSspB | p15A / *cat* / *sspB* | pdCas9-bacteria |
| pGC533-1 | pSspB-rs | p15A / *cat* / *sspB*-rs | pdCas9-bacteria |
| pGC589-1 | pSspB-rs-rbs2 | p15A / *cat* / *sspB*-rs-rbs2 | pGC533-1 |
| pGC591-1 | pSspB-rs-rbs4 | p15A / *cat* / *sspB*-rs-rbs4 | pGC533-1 |
| pGC642-1 | | R6K / *bla kan* / DAS#3(no myc) | pGC015 |
| pGC643-1 | pGFP-DAS#3noMyc | pBR322 / *kan* / GFPmut3b(Δ9)-DAS#3nomyc | pZE27GFP / pGC642-1 |
| pGC702-1 | | R6K / *bla kan* / DAS#3 | pGC642-1 |
| pGC707-1 | pGFP-DAS#3 | pBR322 / *kan* / GFPmut3b(Δ9)-DAS#3 | pGC643-1 |
| pGC799-1 | pClpPX | pSC101* / *kan* / ClpP-ClpX | pZS*24MCS |

**Table S3.** Primers and design details.

| | |
|---|---|
| pGC015 | |
| pKD4 + NheI by Quickchange site directed mutagenesis | |
| GC.013 | GGCCAGTGCCAAGCTTGCTAGCAGATTGCAGCATTACACG |
| GC.014 | CGTGTAATGCTGCAATCTGCTAGCAAGCTTGGCACTGGCC |
| pGC016 | |
| pKD4::DAS#1 (1xmyc-DAS+4-(frt)-KAN-(frt)), by annealing four primers, digesting with ClaI / Nhe I and and ligation into pGC015 digested with the same enzymes | |
| GC.003 | CAAATGATGAAAATTATAGCGAAAATTATGCGGACGCGAGCTAATG |
| GC.004 | CTAGCATTAGCTCGCGTCCGCATAATTTTCGCTATAATTTTCATCATTTGCAGC |
| GC.005 | CGATGAGCAGAAGCTCATCTCGGAAGAGGATCTGGCTG |
| GC.006 | CAGATCCTCTTCCGAGATGAGCTTCTGCTCAT |
| eGC023 | |
| *recA-DAS#1-kan* by recombineering - Template: pGC016 | |
| GC.017 | TAGATGATAGCGAAGGCGTAGCAGAAACTAACGAAGATTTTGAGCAGAAGCTCATCTCGG |
| GC.018 | GGGCCGCAGATGCGACCCTTGTGTATCAAACAAGACGATTAGCTGACATGGGAATTAGCC |
| eGC024 | |
| *ruvB-DAS#1-kan* by recombineering - Template: pGC016 | |
| GC.019 | GGGCGTGGAATCACTTTGGCATAACGCCGCCAGAAATGCCGGAGCAGAAGCTCATCTCGG |
| GC.020 | TTGCCAGTGCCGGATGCGGCGCGAGCGACCAATCCGACTTAGCTGACATGGGAATTAGCC |
| pGC064 | |
| pLAC22::*sspB* by amplification of *sspB* with primers adding BglII (GC.028) or EcoRI (GC.029) sites for cloning into the same sites on pLAC22 | |
| GC.028 | CAGGAAAGATCTATGGATTTGTCACAGCTAACACC |
| GC.029 | ACATGAGAATTCTTACTTCACAACGCGTAATGCC |
| pGC561 | |
| pKD4::DAS#2 - Gene fragment Gibson assembled with PCR amplified pKD4 | |
| Gene fragment | AGTGCAGGTTCTGCAGCCGGTTCTGGTGCAGCTGAACAAAAACTCATCTCTGAAGAAGATCTGGCCGCAAATGATGAAAACTACTCCGAGAATTATGCTGACGCGTCC |

| | |
|---|---|
| Template: pKD4 | |
| GC.467 | CGAGAATTATGCTGACGCGTCCTAATGCTAGCAGATT GCAGCATTACAC |
| GC.468 | GGCTGCAGAACCTGCACTATCGATGCAGGTGGCACTT TTC |
| eGC566 | |
| *recA-DAS#2-kan* by recombineering - Template: pGC561 | |
| GC.469 | TAGATGATAGCGAAGGCGTAGCAGAAACTAACGAAGA TTTTAGTGCAGGTTCTGCAGCC |
| GC.470 | GCCGCAGATGCGACCCTTGTGTATCAAACAAGACGAT TACGGCTGACATGGGAATTAGCC |
| eGC568 | |
| *ruvB-DAS#2-kan* by recombineering - Template: pGC561 | |
| GC.471 | CGGGCGTGGAATCACTTTGGCATAACGCCGCCAGAAA TGCCGAGTGCAGGTTCTGCAGCC |
| GC.472 | GCCAGTGCCGGATGCGGCGCGAGCGACCAATCCGACT TACGGCTGACATGGGAATTAGCC |
| pGC513-1 and pGC533-1 | |
| Both plasmids consist of the pdCas9-bacteria backbone, including the TET promoter and resident RBS. The *sspB* terminator and some flanking sequence are derived from a de novo synthesized chimeric *sspB* construct containing (mf) pdt#3 (Cameron and Collins, 2014 - PMID: 25402616) | |
| pGC467 (intermediate plasmid) | |
| pdCas9-bacteria and Genscript insert assembled by Gibson | |
| pdCas9-bacteria template | |
| GC.360 | CGATGCATATGCCTCACTGACTCGCTACG |
| GC.361 | CGTCACCCATAGATCCTTTCTCCTCTTTAGATC |
| Genscript - pUC57-simple::J23100-B0034-ClpXBind-*SspB-(mf)pdt#3*-B0014 (Jxx and Bxx designations found at http://parts.igem.org/) (intermediate plasmid) (delivered with insert cloned into EcoRV site) | |

| | |
|---|---|
| Insert sequence | GAATTCTTGACGGCTAGCTCAGTCCTAGGTACAGTGC<br>TAGCTACTAGAGAAAGAGGAGAAAAAGCTTATGGGTG<br>ACGACCGTGGTGGTCGTCCGGCTCTGCGTGTTGTTAA<br>AGATACCAGCATCATGAATGATGAAGAGGCATCGGCA<br>GACAACGAAACCGTTATGTCGGTTATTGATGGCGACA<br>AGCCAGATCACGATGATGACACTCATCCTGACGATGA<br>ACCTCCGCAGCCACCAATGGATTTGTCACAGCTAACA<br>CCACGTCGTCCCTATCTGCTGCGTGCATTCTATGAGT<br>GGTTGCTGGATAACCAGCTCACGCCGCACCTGGTGGT<br>GGATGTGACGCTCCCTGGCGTGCAGGTTCCTATGGAA<br>TATGCGCGTGACGGGCAAATCGTACTCAACATTGCGC<br>CGCGTGCTGTCGGCAATCTGGAACTGGCGAATGATGA<br>GGTGCGCTTTAACGCGCGCTTTGGTGGCATTCCGCGT<br>CAGGTTTCTGTGCCGCTGGCTGCCGTGCTGGCTATCT<br>ACGCCCGTGAAAATGGCGCAGGCACGATGTTTGAGCC<br>TGAAGCTGCCTACGATGAAGCGGCGAACAAAAACGAA<br>GAAAACACCAACGAAGTGCCGACCTTTATGCTGAACG<br>CGGGCCAGGCGAACAGAAGACGAGTTTAATAAGGTAC<br>CTCACACTGGCTCACCTTCGGGTGGGCCTTTCTGCGT<br>TTATATACTAGAGAGAGAATATAAAAAGCCAGATTAT<br>TAATCCGGCTTTTTTATTATTTCTCGAGAACCTCCGA<br>TG |
| Template: pUC57-simple::J23100-B0034-ClpXBind-SspB-(mf)pdt#3-B0014 | |
| GC.354 | GAAAGGATCTATGGGTGACGACCGTGGTG |
| GC.355 | CAGTGAGGCATATGCATCGGAGGTTCTCGAG |
| In the case of pGC513-1, the *mf-pdt#3* (Cameron and Collins, 2014 - PMID: 25402616) sequence was removed during Gibson assembly with the wild-type *sspB* | |
| pGC513-1 | |
| pGC467 template | |
| GC.425 | GTGAAGTAAGGTACCTCACACTGGCTC |
| GC.428 | GTGACAAATCCATAGATCCTTTCTCCTCTTTAGATC |
| pGC064 template | |
| GC.427 | AAGGATCTATGGATTTGTCACAGCTAACACCAC |
| GC.426 | GTGTGAGGTACCTTACTTCACAACGCGTAATGCCG |
| In the case of pGC533-1, an intermediate plasmid containing wild-type *sspB* fused to *mf-pdt#3 (*Cameron and Collins, 2014 - PMID: 25402616) was mutagenized by PCR to change *mf-pdt#3* to the wild-type *mf-pdt* sequence (Gur and Sauer, 2008 - PMID: 18852454) | |
| pGC514-1 (Figure S10, intermediate plasmid to pGC533-1, Gibson assembled) | |
| pGC467 template | |
| GC.421 | GTTGTGAAGGCGGCGAACAAAAACGAAG |
| GC.428 | GTGACAAATCCATAGATCCTTTCTCCTCTTTAGATC |
| pGC064 template | |
| GC.427 | AAGGATCTATGGATTTGTCACAGCTAACACCAC |
| GC.420 | GTTCGCCGCCTTCACAACGCGTAATGCCG |

| | |
|---|---|
| pGC533-1 (Gibson assembly to circularize PCR product) | |
| pGC467 template | |
| GC.427 | AAGGATCTATGGATTTGTCACAGCTAACACCAC |
| GC.428 | GTGACAAATCCATAGATCCTTTCTCCTCTTTAGATC |
| pGC595-1 (pSspB-rs-rbs2), pGC596-1 (pSspB-rs-rbs4) were made by "around-the-horn" PCR (Ochman et al., 1988 - PMID: 2852134) with pGC533-1 as template | |
| pGC595-1 | |
| GC.475 | [PHO]TCCTCTTTAGATCTTTTGAATTCTTTTCTC |
| GC.477 | CATCGGATCTATGGATTTGTCACAGC |
| pGC596-1 | |
| GC.475 | [PHO]TCCTCTTTAGATCTTTTGAATTCTTTTCTC |
| GC.479 | GAATGGATCTATGGATTTGTCACAGC |
| eGC658 (Δ*clpX::kan*) | |
| pKD13 template | |
| GC.533 | GTGCGGCACAAAGAACAAAGAAGAGGTTTTGACCCATGATTCCGGGGATCCGTCGACC |
| GC.534 | CCCTTTTTGGTTAACTAATTGTATGGGAATGGTTAATTATGTAGGCTGGAGCTGCTTCG |
| pGC642 (intermediate plasmid of DAS#3 lacking myc tag) | |
| pGC015 template | |
| GC.520 | CTATGCGGACGCCTCCTAATGCTAGCAGATTGCAGC |
| GC.521 | CTCACCATCGCCTTCACCATCGATGCAGGTGGCACTTTTC |
| gblock (inserted by Gibson) | |
| GC.523 | GGTGAAGGCGATGGTGAGGGAGATGGTGAGGGTGATGCGGAAGCAGCGGCAAAGGAGGCTGCAGCAAAAGAGGCCGCGGCCAAAGAGGCAGCAGCAAAAGCCTTAGAGGCAGAGGCAGCCGCTAAGGAAGCCGCCGCGAAGGAAGCCGCAGCTAAAGCAGCTGCAAATGATGAAAATTACTCGGAGAACTATGCGGACGCCTCC |
| pGC702 (pKD4::DAS#3) | |
| pGC642 template, product circularized by Gibson | |
| GC.615 | CTCATCTCGGAAGAGGATCTGGCTGCAAATGATGAAAATTACTCGGAGAAC |
| GC.616 | GATCCTCTTCCGAGATGAGCTTCTGCTCTGCTTTAGCTGCGGCTTCCTTC |
| eGC698 | |
| *recA-DAS#3-kan* by recombineering - Template: pGC702 | |
| GC.611 | GTAGATGATAGCGAAGGCGTAGCAGAAACTAACGAAGATTTTGGTGAAGGCGATGGTGAG |

| | |
|---|---|
| GC.612 | GGCCGCAGATGCGACCCTTGTGTATCAAACAAGACGA TTAGGCTGACATGGGAATTAGCC |
| eGC699 | |
| *ruvB-DAS#3-kan* by recombineering - Template: pGC702 | |
| GC.613 | CGGGCGTGGAATCACTTTGGCATAACGCCGCCAGAAA TGCCGGGTGAAGGCGATGGTGAG |
| GC.614 | TGCCAGTGCCGGATGCGGCGCGAGCGACCAATCCGAC TTAGGCTGACATGGGAATTAGCC |
| eGC706 (ΔsspB lacZ-DAS#3) | |
| pGC702 template | |
| GC.541 | AGCGCCGGTCGCTACCATTACCAGTTGGTCTGGTGTC AAAAAGGTGAAGGCGATGGTGAG |
| GC.542 | CGCGAAATACGGGCAGACATGGCCTGCCCGGTTATTA TTAGGCTGACATGGGAATTAGCC |
| pZE27GFP degron tags (figure S3) were all constructed as IDT gBlocks Gibson assembled to the same pZE27GFP (Cameron and Collins, 2014 - PMID: 25402616) PCR product | |
| pZE27GFP template (linearization removed C-terminal 9AA, GITHGMDELYK) | |
| GC.518 | GAACTATGCGGACGCCTCCTAATAAGCTTGATGGGGG ATC |
| GC.519 | CTCACCATCGCCTTCACCAATCCCAGCAGCTGTTACA AAC |
| gblock (inserted by Gibson) | |
| pGC639 (GFP-GE-FL-DAS+4) | |
| GC.522 | GGTGAAGGCGATGGTGAGGGGGACGGAGAGGGTGACG GCGGGGGTAGTGGTGGCGGGGGGTCAGGAGGGGGTAG TGCCGCTAACGATGAAAATTATAGCGAGAACTATGCG GACGCCTCC |
| pGC640 (GFP-GE-RLP-DAS+4) | |
| GC.524 | GGTGAAGGCGATGGTGAGGGTGATGGGGAGGGTGACC CGGCCCCTGCTCCCGCTCCTGCACCTGCCCCAGCGCC AGCTCCGGCACCGGCACCTGCTGCAGCCAATGATGAG AATTACAGTGAGAACTATGCGGACGCCTCC |
| Figure S9 | |
| pGC643 (GFP-GE-RLA-DAS+4) | |
| GC.523 | GGTGAAGGCGATGGTGAGGGAGATGGTGAGGGTGATG CGGAAGCAGCGGCAAAGGAGGCTGCAGCAAAAGAGGC CGCGGCCAAAGAGGCAGCAGCAAAAGCCTTAGAGGCA GAGGCAGCCGCTAAGGAAGCCGCCGCGAAGGAAGCCG CAGCTAAAGCAGCTGCAAATGATGAAAATTACTCGGA GAACTATGCGGACGCCTCC |
| pGC707 (GFP-DAS#3) Template pGC643, recircularization by Gibson | |
| GC.615 | CTCATCTCGGAAGAGGATCTGGCTGCAAATGATGAAA ATTACTCGGAGAAC |

| | |
|---|---|
| GC.616 | GATCCTCTTCCGAGATGAGCTTCTGCTCTGCTTTAGC TGCGGCTTCCTTC |
| Figure S1 | |
| pGC332 (pCAS9-CR4::sspB-ssrA) - By Gibson Assembly | |
| Template pCas9-CR4 | |
| GC.197 | GGCATTACGCGTTGTGAAGGCTGCTAACGACGAAAAC TAC |
| GC.196 | GGTGTTAGCTGTGACAAATCCATAGATCCGAAGTCCT CTTTAGATC |
| Template pGC064 | |
| GC.194 | ATGGATTTGTCACAGCTAACACC |
| GC.195 | CTTCACAACGCGTAATGCC |
| pGC334 (pCAS9-CR4::sspB-LAA) - By Gibson Assembly | |
| GC.198 | GGCATTACGCGTTGTGAAGGCTCTGGCTGCTTAACTC |
| GC.196 | GGTGTTAGCTGTGACAAATCCATAGATCCGAAGTCCT CTTTAGATC |
| Template pGC064 | |
| GC.194 | ATGGATTTGTCACAGCTAACACC |
| GC.195 | CTTCACAACGCGTAATGCC |
| pGC336 (pCAS9-CR4::sspB) - By Gibson Assembly | |
| GC.199 | GGCATTACGCGTTGTGAAGTAACTCGAGTAAGGATCT CCAG |
| GC.196 | GGTGTTAGCTGTGACAAATCCATAGATCCGAAGTCCT CTTTAGATC |
| Template pGC064 | |
| GC.194 | ATGGATTTGTCACAGCTAACACC |
| GC.195 | CTTCACAACGCGTAATGCC |
| Figure S2 | |
| pGC779 (pClpP/X) - By Gibson Assembly | |
| Template pZS*24MCS | |
| GC.692 | CTGGTGAATAAGCTTGATATCGAATTCCTGCAGC |
| GC.693 | GCTTTTCGCACGCTCGAGTCGACAGTTCATAGG |
| Template MG1655 gDNA | |
| GC.694 | CGACTCGAGCGTGCGAAAAGCCTCTTTCGG |
| GC.695 | CGATATCAAGCTTATTCACCAGATGCCTGTTGCG |
| dnaA-DAS#2-kan | |
| Template pGC561-1 | |
| GC.494 | GAGCCACGATATCAAAGAAGATTTTTCAAATTTAATC AGAACATTGTCATCGAGTGCAGGTTCTGCAGCC |
| GC.495 | TAGCGGTTTTAATAAATGCTCACGTTCTACGGTAAAT TTCATAGGTTTACGGCTGACATGGGAATTAGCC |
| dnaA-DAS#3-kan | |

| | |
|---|---|
| Template pGC642 | |
| GC.587 | ATCAAAGAAGATTTTTCAAATTTAATCAGAACATTGT CATCGGGTGAAGGCGATGGTGAG |
| GC.588 | TAATAAATGCTCACGTTCTACGGTAAATTTCATAGGT TTAGGCTGACATGGGAATTAGCC |
| dnaG-DAS#2-kan | |
| Template pGC561-1 | |
| GC.502 | CAACGAAGAACGCCTGGAGCTCTGGACATTAAACCAG GAGCTGGCGAAAAAGAGTGCAGGTTCTGCAGCC |
| GC.503 | GCTGTCGGGGGCTTCCCGATCGCTCTTCGGCACTTAA GCCGTTAAATCACGGCTGACATGGGAATTAGCC |
| holC-DAS#2-kan | |
| Template pGC561-1 | |
| GC.593 | TACCGCGTGGCTGGTTTCAACCTGAATACGGCAACCT GGAAAGGTGAAGGCGATGGTGAG |
| GC.594 | GGCTGTTCGATATCTTGTGGGTTATATGTCTTTTCCA TTAGGCTGACATGGGAATTAGCC |
| holD-DAS#2-kan | |
| Template pGC561-1 | |
| GC.609 | CAAATTTGCACATATGAACACGATTTCTTCCCTCGAA ACGACGGTGAAGGCGATGGTGAG |
| GC.610 | GTGGGCGCGTTGTTCAATGTGGTAAGCCGCCGGTAAA TCAGGCTGACATGGGAATTAGCC |
| ligA-DAS#2-kan | |
| Template pGC561-1 | |
| GC.482 | ACTGGGCATTGAAGTCATCGACGAAGCGGAAATGCTG CGTTTGCTGGGTAGCAGTGCAGGTTCTGCAGCC |
| GC.483 | CGGCATTATCGTATTGGCTATTTCAATCAGCTGCTCT TTTTCCATCTCACGGCTGACATGGGAATTAGCC |
| ligA-DAS#3-kan | |
| Template pGC642 | |
| GC.597 | GAAGTCATCGACGAAGCGGAAATGCTGCGTTTGCTGG GTAGCGGTGAAGGCGATGGTGAG |
| GC.598 | CGTATTGGCTATTTCAATCAGCTGCTCTTTTTCCATC TCAGGCTGACATGGGAATTAGCC |
| polA-DAS#2-kan | |
| Template pGC561-1 | |
| GC.484 | TGTGCCGTTGCTGGTGGAAGTGGGGAGTGGCGAAAAC TGGGATCAGGCGCACAGTGCAGGTTCTGCAGCC |
| GC.485 | CAGCTTATGTTGCTTACTTACGAAAAAAGGCATGTTC AGGCGAATCTTACGGCTGACATGGGAATTAGCC |

| | |
|---|---|
| polA-DAS#3-kan | |
| Template pGC642 | |
| GC.543 | CTGGTGGAAGTGGGGAGTGGCGAAAACTGGGATCAGG CGCACGGTGAAGGCGATGGTGAG |
| GC.544 | TTGCTTACTTACGAAAAAAGGCATGTTCAGGCGAATC TTAGGCTGACATGGGAATTAGCC |
| priA-DAS#2-kan | |
| Template pGC561-1 | |
| GC.486 | ACCGGATTCCCGTAAGGTGAAATGGGTGCTGGATGTT GATCCGATTGAGGGTAGTGCAGGTTCTGCAGCC |
| GC.487 | TGTGATGAATATTGAATTTTTCGATCCGCCTCGCATC GTGAGCGGTTTACGGCTGACATGGGAATTAGCC |
| recB-DAS#2-kan | |
| Template pGC561-1 | |
| GC.490 | GTTGATTGCCCTGATGGATGAGATGTTTGCCGGTATG ACCCTGGAGGAGGCGAGTGCAGGTTCTGCAGCC |
| GC.491 | AGCTGTTTGTGCTCCACAGCTTCCAGTAATTGCTTTT GCAATTTCATTACGGCTGACATGGGAATTAGCC |
| recB-DAS#3-kan | |
| Template pGC642 | |
| GC.545 | AACGCCGGGTTGATTGCCCTGATGGATGAGATGTTTG CCGGTGGTGAAGGCGATGGTGAG |
| GC.546 | TGCTCCACAGCTTCCAGTAATTGCTTTTGCAATTTCA TTAGGCTGACATGGGAATTAGCC |
| recC-DAS#2-kan | |
| Template pGC561-1 | |
| GC.492 | CGTTGAACAGTCGCAACGTTTCCTGTTACCGCTGTTT CGCTTTAATCAGTCAAGTGCAGGTTCTGCAGCC |
| GC.493 | CGCATCATAAAGTAAGCGGATAGATTGCGCAATTTTT ATACAGCACTCACGGCTGACATGGGAATTAGCC |
| recC-DAS#3-kan | |
| Template pGC642 | |
| GC.547 | TCGCAACGTTTCCTGTTACCGCTGTTTCGCTTTAATC AGTCAGGTGAAGGCGATGGTGAG |
| GC.548 | AAGTAAGCGGATAGATTGCGCAATTTTTATACAGCAC TCAGGCTGACATGGGAATTAGCC |
| rnhA-DAS#3-kan | |
| Template pGC642 | |
| GC.580 | ATGAATCCCACACTGGAAGATACAGGCTACCAAGTTG AAGTTGGTGAAGGCGATGGTGAG |
| GC.581 | CGGTTGGAGCCACCCGGCAATGTCGTAAACCACAGGC TTAGGCTGACATGGGAATTAGCC |

| | |
|---|---|
| Figure S10 | |
| Integration of pGC533-1 fragment into the genomic copy of *sspB* | |
| Template pGC533-1 | |
| GC.440 | GTCAGGATGAGTGTCATCATCGTGATCTGGCTTGTCG CATATGCATCGGAGGTTCTCGAG |
| GC.441 | GTCCCTATCTGCTGCGTGCATTCTATGAGTGGTGTTG TTACTAGTGCTTGGATTCTCACC |

**References**

1. R. Lutz, H. Bujard, Independent and tight regulation of transcriptional units in Escherichia coli via the LacR/O, the TetR/O and AraC/I1-I2 regulatory elements. *Nucleic Acids Res* **25**, 1203-1210 (1997).
2. T. A. Bell, T. A. Baker, R. T. Sauer, Interactions between a subset of substrate side chains and AAA+ motor pore loops determine grip during protein unfolding. *eLife* **8** (2019).
3. X. Chen, J. L. Zaro, W. C. Shen, Fusion protein linkers: property, design and functionality. *Adv Drug Deliv Rev* **65**, 1357-1369 (2013).
4. K. E. McGinness, T. A. Baker, R. T. Sauer, Engineering controllable protein degradation. *Mol Cell* **22**, 701-707 (2006).
5. K. A. Datsenko, B. L. Wanner, One-step inactivation of chromosomal genes in Escherichia coli K-12 using PCR products. *Proceedings of the National Academy of Sciences of the United States of America* **97**, 6640-6645 (2000).
6. D. E. Cameron, J. J. Collins, Tunable protein degradation in bacteria. *Nature biotechnology* **32**, 1276-1281 (2014).
